# Expansion of nitrogenase-like enzymes involved in microbial assimilation of volatile organic sulfur compounds

**DOI:** 10.64898/2026.08.20.746119

**Authors:** Nicole L. Márquez Reyes, Ana Arroyo Carriedo, Justin A. North, Kathryn R. Fixen

## Abstract

Organosulfur compounds are the predominant source of sulfur in terrestrial environments, and bacteria in these environments require enzymes that allow them to assimilate organosulfur compounds. Most described enzymes involved in organosulfur assimilation require oxygen, and understanding of enzymes that function under anoxic conditions is limited. Recently, the enzyme methylthio-alkane reductase (Mar), a nitrogenase-like enzyme that reduces the volatile organic sulfur compounds (VOSCs) methylthio ethanol (MT-EtOH), dimethyl sulfide (DMS), and ethyl methyl sulfide (EMS) in the absence of oxygen was identified in the purple nonsulfur bacterium *Rhodospirillum rubrum.* However, another purple nonsulfur bacterium, *Rhodopseudomonas palustris,* has three loci containing nitrogen fixation-like (NFL) genes with high sequence similarity to Mar, suggesting there are other Mar-like enzymes that may serve distinct roles. Here, we tested if these NFL genes in *R. palustris* are required to assimilate VOSCs. Transcriptomic sequencing (RNA-seq) analysis revealed that all NFL genes in *R. palustris* are up-regulated in response to sulfur limitation, supporting a role in sulfur assimilation. Only disruption of the NFL genes encoded by RPA2634*-*37, renamed *marBHDK1*, resulted in fitness defects when EMS, DMS, and dimethylsulfoniopropionate (DMSP) were provided as a sulfur source, indicating it is a functional Mar enzyme. The NFL genes encoded by RPA2347-48 and RPA2353-54, renamed *marKD2* and *marHB2*, respectively, were required for activity when MT-EtOH or ethanedithiol was provided as a sulfur source but not DMS, EMS, or DMSP. No activity was observed with the third NFL loci that includes RPA2363 and RPA2364, renamed *nflDK*. Overall, the results demonstrate that two homologs of Mar in *R. palustris* are capable of VOSC reduction, with one specialized for simple VOSCs and the other acting preferentially on a substrate with an additional functional group.

**IMPORTANCE:** VOSCs in freshwater environments play a role in atmospheric processes, impacting global weather patterns. Bacteria are central to cycling sulfur in these environments, driving sulfur transformations even in oxygen-limited environments where sulfate is scarce but organosulfur compounds are abundant. While many oxygen-dependent reactions have been described that contribute to VOSC cycling, anaerobic pathways remain much less understood. Mar enzymes represent a newly characterized mechanism for anoxic VOSC assimilation. Here we find that two Mar homologs in *R. palustris* are the result of functional specialization of different Mar isozymes. Studying their activity expands our understanding of microbial strategies for sulfur turnover and sheds light on anaerobic sulfur metabolism.

## INTRODUCTION

Sulfur is an essential element for life, as it is an important component of coenzymes and the amino acids cysteine and methionine. Sulfate is most abundant in marine ecosystems but is much lower in abundance in freshwater ecosystems, often present at concentrations below 100 pM (1–3). Volatile organic sulfur compounds (VOSCs) in freshwater environments have been studied because of their role in atmospheric processes such as acid rain and cloud formation and as a source of odorous emissions in drinking water and wastewater treatment (4). With increasing anthropogenic emissions of VOSCs, understanding degradation pathways is essential for assessing how microbial processes may mitigate their release. However, the assimilation of VOSCs, particularly under anoxic conditions, remains poorly characterized and is currently understood mainly through studies of dimethyl sulfide (DMS) methyl Htransfer activity in methanogenic archaea (5–8).

More recently, a nitrogenase-like enzyme capable of reducing the C-S bond in VOSCs has been identified in the purple nonsulfur bacterium, *Rhodospirillum rubrum*. Under sulfur-limiting conditions, this enzyme, known as *m*ethylthio-*a*lkane *r*eductase (Mar), can reduce DMS, 2-methylthio ethanol (MT-EtOH), and ethyl methyl sulfide (EMS) to generate methanethiol (MT), which is incorporated via the methionine synthesis cycle (2, 3). Mar activity also results in the release of gaseous products during reduction of VOSCs. Ethylene (C_2_H_4_), an important plant hormone (9, 10) and a precursor for plastic production, is released when Mar reduces MT-EtOH (3), and may account for the ethylene released by waterlogged soil (11). Mar reduction of DMS and EMS results in the release of the potent greenhouse gas, methane (CH_4_), and ethane (C_2_H_6_), respectively (3). This finding indicates that Mar activity plays a role in turnover of a variety of VOSCs under anoxic conditions, and understanding its regulation and activity will help understand its role in terrestrial environments.

Mar enzymes are part of the nitrogenase enzyme superfamily (3). Nitrogenase is the key enzyme involved in nitrogen fixation, reducing atmospheric nitrogen to produce ammonia (NH_3_) and hydrogen (H_2_) (12–16). Mar enzymes are composed of homologous components to those found in nitrogenase. Like NifH, MarH forms a homodimeric complex in which conserved cysteines coordinate a 4Fe-4S cluster, enabling it to function as an ATPHdependent reductase (17, 18). MarDK is homologous to NifDK, and forms a heterotetrameric complex that also has six conserved cysteines that coordinate a P-cluster-like cofactor (**Fig. 1**)(3, 13). However, unlike NifDK, MarDK does not bind an iron molybdenum cofactor and binds a putative [Fe_8_S_9_C]-cluster, similar to the L-cluster produced by NifB, which is involved in cofactor biosynthesis for nitrogenase (17, 18) (**Fig. 1**). This L-like cluster is made by MarB, which is in an operon with MarHDK, and is homologous to NifB (**Fig. 1**)(3).

**Figure 1.**
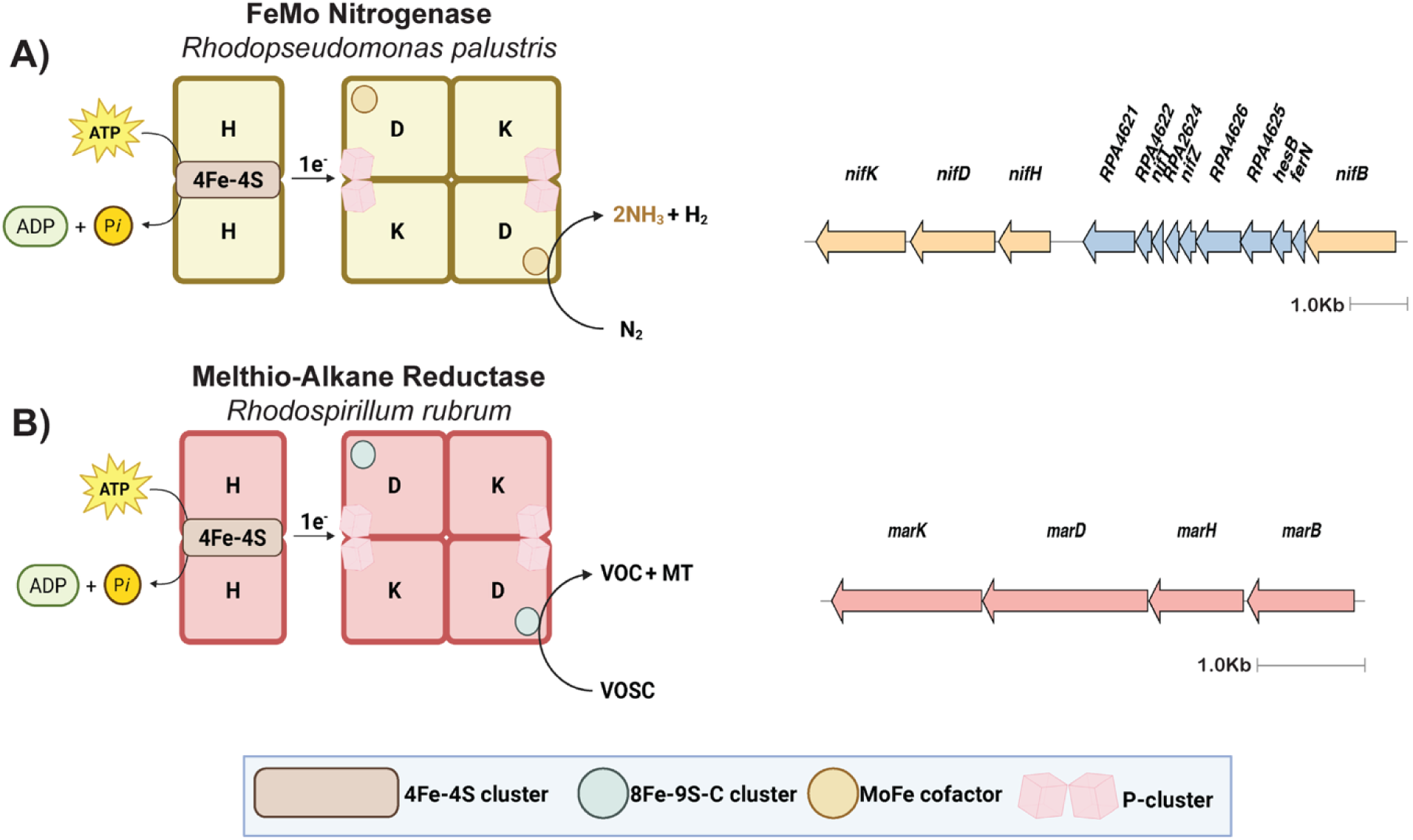
Simplified schematic and gene neighborhoods of **(A)** FeMo nitrogenase components (16) and its respective gene cluster in *R. palustris.* **(B)** MarHDK of *R. rubrum* and the operon that encodes it. In the reaction VOSC = volatile organic sulfur compound, VOC = volatile organic compound, and MT = methanethiol. NifB and MarB, shown in the gene neighborhoods, are the respective maturases for the FeMo cofactor and the 8Fe-9S-C (”mar2” (17)) cluster.

The purple nonsulfur bacterium *Rhodopseudomonas palustris* thrives in freshwater sediments where sulfate availability is likely to be limited, conditions under which the ability to access VOSCs as alternative sulfur sources would be advantageous. Notably, *R. palustris* encodes nitrogen fixation-like (NFL) genes that include two MarBH and three MarDK homologs that cluster phylogenetically with the MarBHDK proteins of *R. rubrum* (3). It is unclear if these NFL genes in *R. palustris* play a role in assimilating VOSCs, and if the different homologs represent functional diversity in this group. The purpose of this study was to gain insight into the function of the NFL genes in *R. palustris.* Using RNA-seq, we found all NFL genes are upregulated when *R. palustris* is grown under sulfate-limiting conditions, supporting a role in sulfur assimilation. We also found that only the NFL genes RPA2634*-*37, which encode the homolog we refer to as Mar1, are required to assimilate the VOSCs DMSP, DMS, and EMS. However, a second loci containing NFL genes RPA2347-48 and RPA2353-54, which encode the homolog we refer to as Mar2, contributes to ethylene production when *R. palustris* is grown with MT-EtOH or ethanedithiol as a sulfur source, indicating this Mar homolog is not involved in reducing simple VOSCs and has more activity with larger VOSCs containing different functional groups. Together, these data indicate that at least two of the three NFL gene clusters in *R. palustris* play a role in assimilation of VOSCs as a sulfur source and have distinct substrate preferences, expanding the VOSCs *R. palustris* can access as a sulfur source.

## RESULTS

### R. palustris *CGA009* encodes multiple NFL genes homologous to Mar and can grow on VOSCs

Similar to *R. rubrum,* the *R. palustris* gene cluster RPA2634*-*37, referred to as Mar1, includes the genes *marBHDK1*. The MarHDK1 has the highest percent identity with MarHDK from *R. rubrum* (**Fig. 2**)(3). *R. palustris* also has one other set of NFL genes, RPA2353 and RPA2354, that encode homologs of MarB and MarH and are separated by 4.5 kb from the NFL genes RPA2347-48, which encode homologs of MarD and MarK. This group of NFL genes will be referred to as Mar2 and include the genes *marBHDK2* (**Fig. 2**). A third homolog of MarDK is encoded by the NFL genes RPA2363 and RPA2364 and are 9 kb downstream of the MarH homolog in Mar2 (**Fig. 2**). This gene cluster will be referred to as NflDK3.

**Figure 2.**
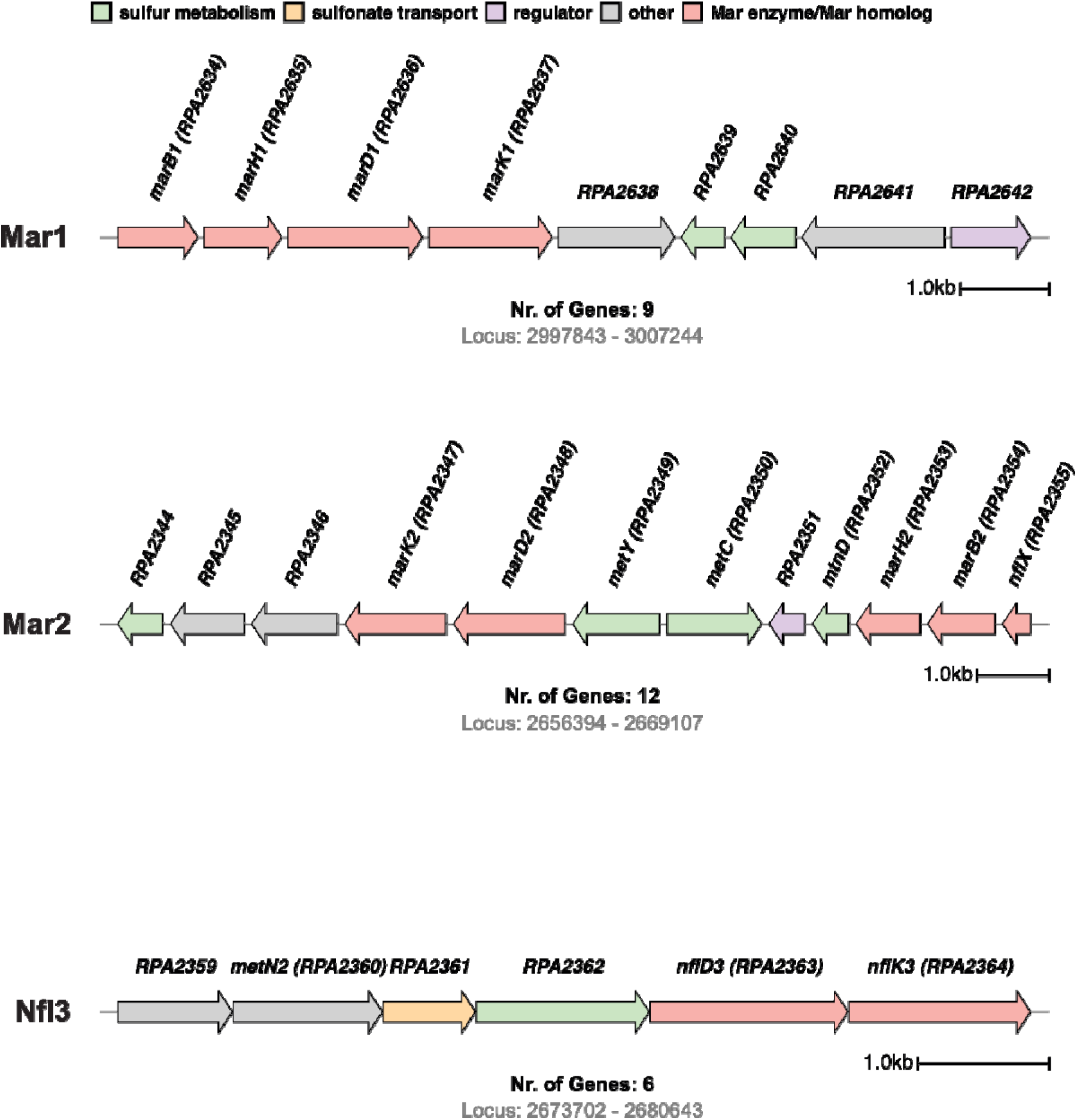
Mar homologs in *Rhodopseudomonas palustris.* *R. palustris* encodes three distinct. Mar enzymes. These are found in three gene neighborhoods (Mar1, Mar2, and Nfl3) with other sulfur metabolism and transport genes. Color legend indicates gene function: [green = sulfur metabolism, yellow = sulfonate transport, purple = regulation, grey = other, and red = Mar enzyme/Mar homolog].

Mar1, Mar2, and NflDK3 are surrounded by other genes involved in sulfur assimilation a**nd** transport (**Fig. 2**). In Mar2, *marB2* and *marH2* are flanked by *mtnD,* which encodes **an** acidoreductone dioxygenase involved in methionine salvage, and RPA2355, a homolog of N**ifX** (**Fig. 2**). NifX binds and stabilizes the cofactor generated by NifB (19). Genes annotated **as** *metY* (RPA2349) and *metC* (RPA2350) are encoded next to *marD2* (**Fig. 2**). The genes encoding NflDK3 are adjacent to genes encoding a putative methionine transporter and O-acetylhomoserine sulfhydrylase (**Fig. 2**). The placement of the NFL genes encoding Mar1, Mar2, and NflDK3 adjacent to or even in a potential operon with other sulfur metabolism genes indicate that these homologs likely participate in sulfur metabolism.

To determine if the NFL genes encoding Mar1, Mar2, and NflDK3 are likely to bind metallocofactors like MarHDK from *R. rubrum*, we determined if residues critical for binding the metallocofactors in MarHDK are present in MarHDK1, MarHDK2, and NflDK3. MarH coordinates a single [4Fe-4S] cluster with cysteines at position 97 and 133 (*R. rubrum* numbering) (17, 18). Both MarH1 and MarH2 in *R. palustris* have an ATP-binding site and the cysteine residues required for coordination of the [4Fe-4S] cluster (**Fig. S1**), and the amino acid identity between MarH1 and MarH2 are highly similar, sharing around 94% amino acid identity (**Fig. 3B**), indicating both MarH homologs in *R. palustris* likely act as an ATP-dependent reductase.

**Figure 3.**
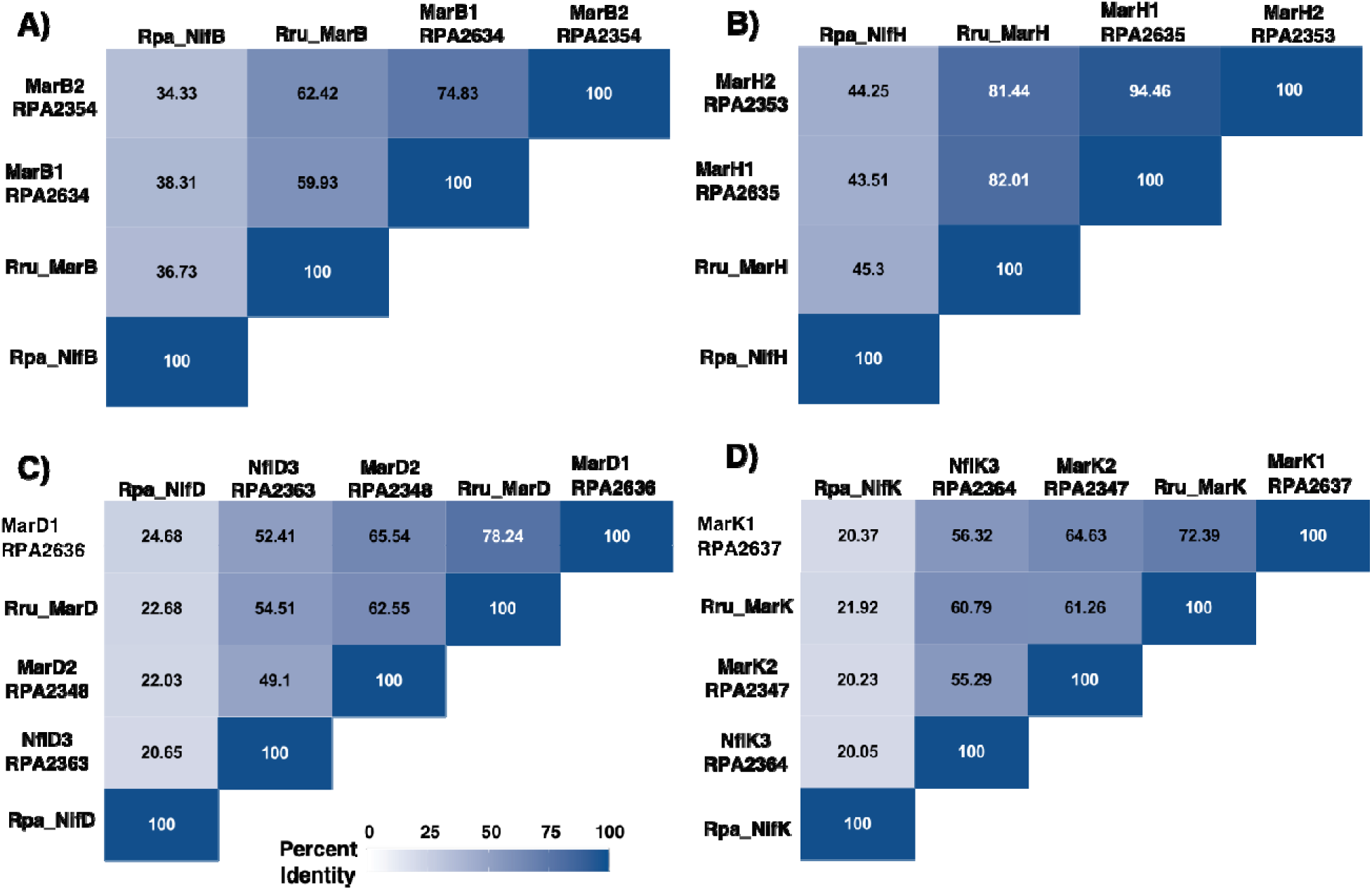
Precent identity Comparison of Mar enzymes and nitrogenase. **(A)** Precent Identity matrix of MarB homologs from *R.palustris* and *R. rubrum* and NifB from *R. palustris*. **(B)** Precent matrix of MarH homologs from *R. palustris* and *R. rubrum*, and NifD from *R. palustris*. **(C)** Percent identity matrix of MarD homologs from *R. palustris* and *R. rubrum*, and NifD from *R. palustris*. **(D)** Percent identity matrix of Mark homologs from *R. palustris* and *R. rubrum*, and NiFK from *R. palustris*.

The cysteine residues in *R. rubrum* MarD and MarK that coordinate a P-cluster are conserved in all of the NFL sequences with similarity to MarD and MarK in *R. palustris,* indicating the potential to bind a P-cluster (3, 17, 18)(“#” in **Fig. S1**). The MarD homologs all share the cysteine at position 270 and the histidine at position 429 in *R. rubrum* that coordinates an L-like cluster. This suggests that all three *R. palustris* NFL sequences with similarity to MarD also have the potential to bind a metallocofactor structurally similar to the L-cluster. Based on percent identity of the Mar homologs in *R. palustris* compared to Mar in *R. rubrum*, MarHDK1 is most similar to MarHDK from *R. rubrum*, further suggesting MarHDK1 has similar activity to MarHDK from *R. rubrum* (**Fig. 3**). The MarD and MarK homologs from Mar1 and Mar2 share ∼65% amino acid identity (**Fig. 3C-D**), whereas MarDK1 and NflDK3 are more dispara**te,** sharing ∼52-56% amino acid identity (**Fig. 3C-D**).

Synthesis of the putative L-like cluster bound by MarDK requires MarB, the NifB homolog fou**nd** in the *marBHDK* operon in *R. rubrum.* MarB is similar to NifB in that it contains many of t**he** same domains as NifB, but MarB lacks the NifX-like domain contained in some NifB (20). In ***R.*** *palustris,* MarB homologs found in Mar1 and Mar2 share 74.8% amino acid identity and exhi**bit** all conserved motifs found in MarB of *R. rubrum* (**Fig. 3A; Fig. S1**). While MarHDK1 in ***R.*** *palustris* shows the highest similarity to MarHDK from *R. rubrum*, MarB1 is slightly less simi**lar** to MarB from *R. rubrum*, sharing only 59.9% amino acid identity, whereas MarB2 shares 65.5**%** amino acid identity with MarB from *R. rubrum*. Together this suggests that the two MarB homologs in *R. palustris* could play a role in biosynthesis of an L-like-cluster.

These results indicate that *R. palustris* encodes at least one enzyme functionally similar to MarHDK from *R. rubrum*. To determine if *R. palustris* possesses a functional Mar system, it was cultivated in defined medium with VOSCs provided as the sole sulfur source. In defined medium lacking an added sulfur source, *R. palustris* was unable to reach high cell densities, but growth was restored when the Mar substrates MT-EtOH, DMS, or EMS were provided (**Fig. 4**). *R. palustris* was unable to use these compounds as a carbon source in either sulfate-replete or sulfate-limited medium (data not shown), indicating that they are used specifically as sulfur sources. Growth was also observed with DMSP, which is predicted to be converted to DMS, as *R. palustris* encodes a putative DMSP lyase (RPA2344) (**Fig. 4**). Collectively, these data indicate that there is a functional Mar system in *R. palustris*.

**Figure 4.**
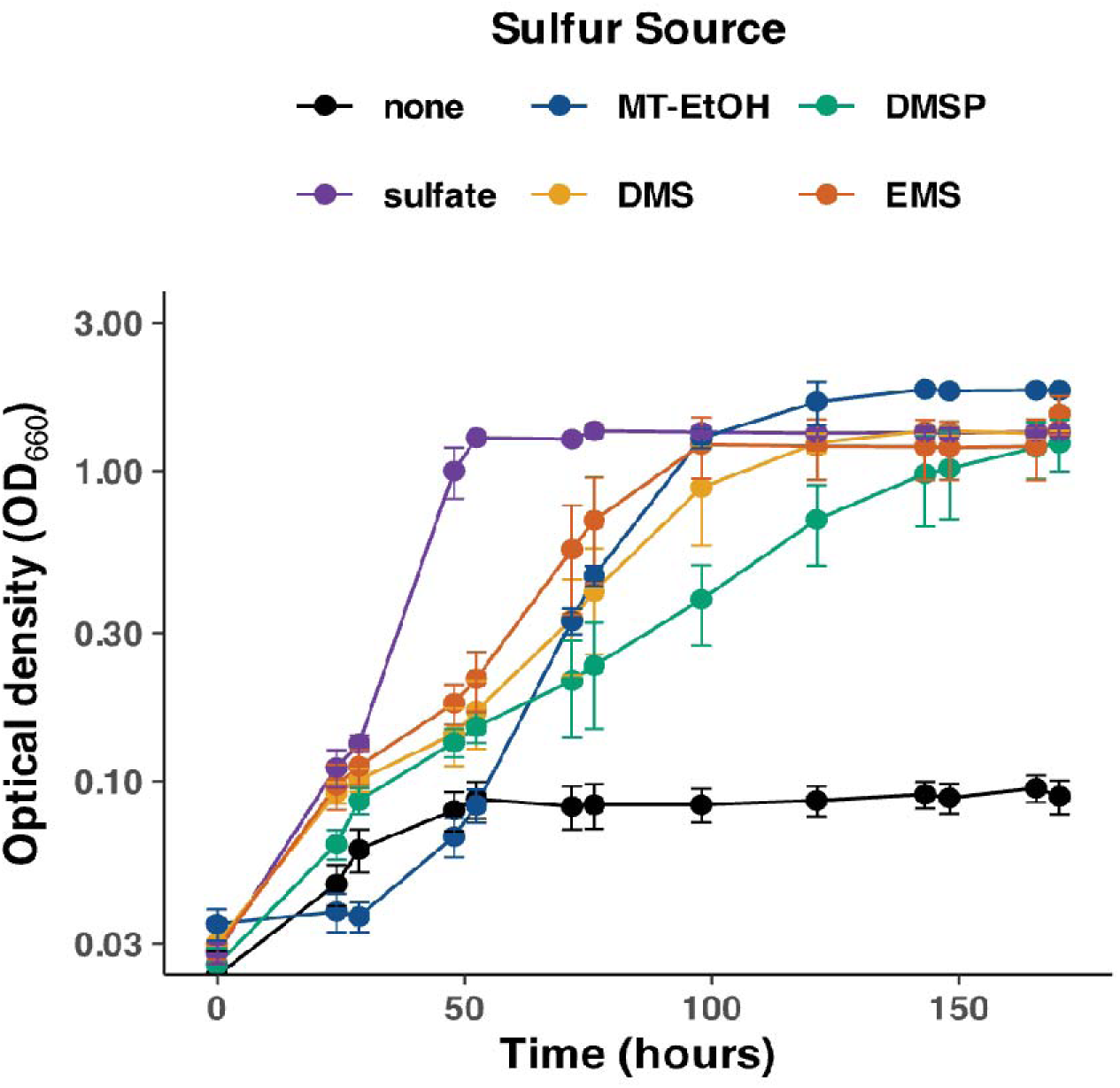
*R. palustris* CGA009 assimilates methylthio ethanol (MT-EtOH), dimethyl sulfide (DMS), dimethylsulfoniopropionate (DMSP), and ethyl-methyl sulfide (EMS). Growth was measured using optical density at 660 nm over time. Cultures with sulfate, DMS, DMSP, and EMS provided with 20 mM acetate as a carbon source, and MT-EtOH was provided 10 mM malate as a carbon source. All VOSCs were added in the final concentration of 1 mM. Error bars represent the standard deviation of three biological replicates.

### All Mar homologs are up-regulated under sulfate-limiting conditions

In sulfate limited conditions, bacteria upregulate pathways that enable them to scavenge remaini**ng** sulfate and assimilate alternative sulfur sources (21). If a NFL gene is required for sul**fur** assimilation, we anticipated that expression of the NFL gene would increase under sulfate**-** limited conditions. Using RNA-seq, gene expression changes in *R. palustris* shifted from sulfate**-** replete to sulfate-limiting conditions or grown with MT-EtOH as the sulfur source versus grow**th** with sulfate as the sulfur source was determined. In all cases, *R. palustris* was grow**n** photoheterotrophically with 10 mM malate supplied as the carbon source.

Under these conditions, gene expression patterns indicated that *R. palustris* was experiencing sulfate limitation. We observed increased expression of a gene cluster involved in sulfate transport and activation (RPA0749-53) (**Fig. S2; Supplemental Dataset 1**). Genes associated with amino acid transport were also upregulated, including those encoding a putative methionine transporter (RPA1426-1429) and a general amino acid transporter (RPA2628-33), both of which showed higher expression under sulfate limiting conditions and when MT EtOH was supplied as the sulfur source (**Fig. S2; Supplemental Dataset 1**). Similarly, genes required for the assimilation of other organic sulfur compounds, such as sulfonates, were upregulated. This included an aliphatic sulfonate transporter and associated assimilation enzymes, including the nitrogenase like enzyme, isethionate reductase, IsrHDK (RPA2608-18) (**Fig. S2; Supplemental Dataset 1**). As shown in **Fig. 5B**, all NFL genes in Marl, Mar2, and NflDK3 in *R. palustris* are up-regulated when cultures were shifted to sulfate-limiting conditions or when MT-EtOH is provided as the sole sulfur source. These data indicate that in response to sulfur limitation, *R. palustris* increases expression of several potential pathways for accessing alternative sulfur-containing compounds, including all of the NFL genes in Marl, Mar2, and NflDK3.

**Figure 5.**
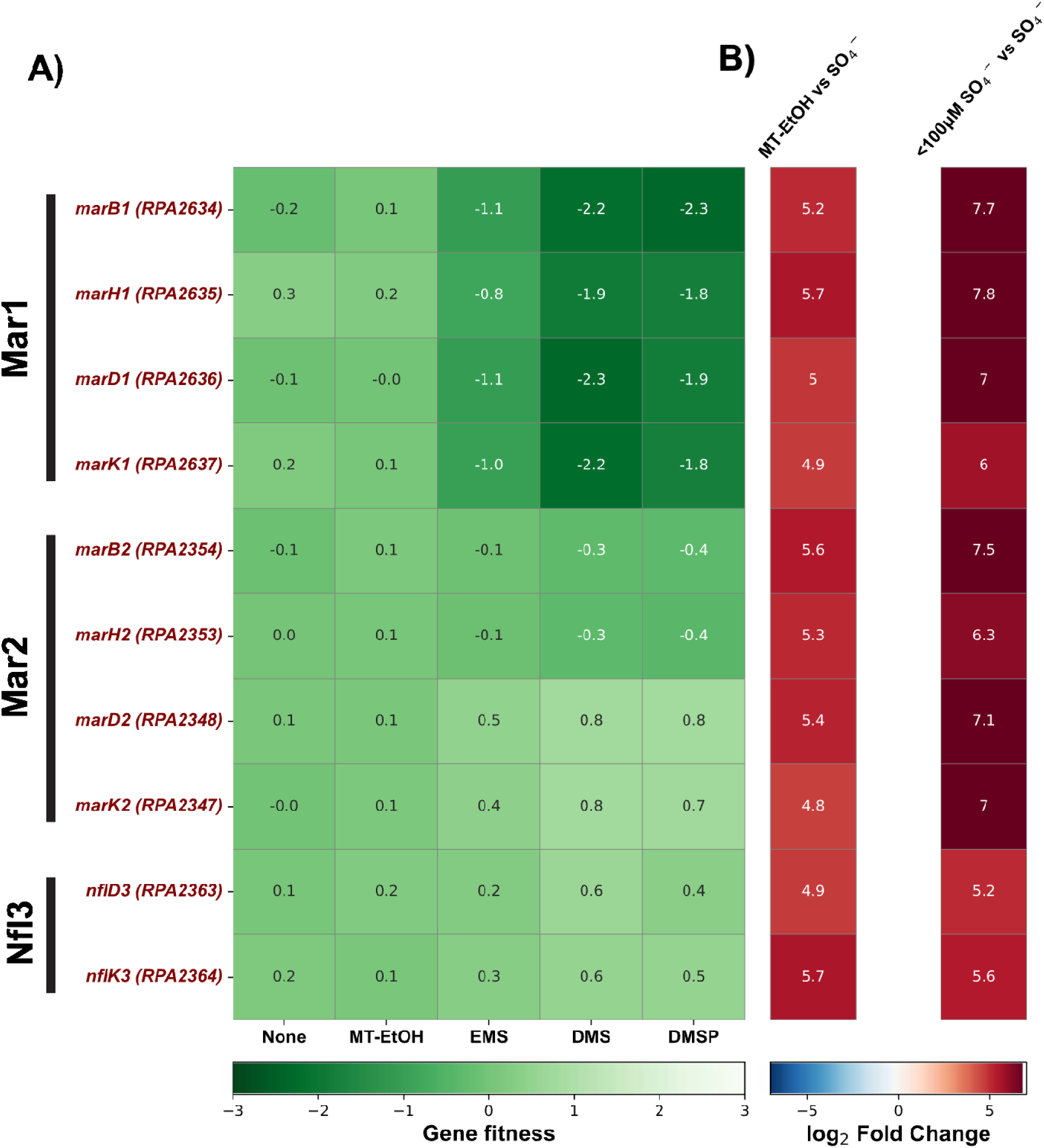
All Mar homologs are up-regulated in sulfate-limiting conditions, but only MarBHDK1 (RPA2634-37) is required for assimilation of DMS, DMSP, and EMS. **(A)** Heatmap of average gene fitness values for each Mar homolog when either DMS, DMSP, EMS or MT-EtOH was provided as a sulfur source. Gene fitness values are bounded by −3 to 3. **(B)** Log_2_fold change in gene expression of genes encoding Mar homologs in sulfate-limited medium (<100 μM SO_4_^-^) versus sulfate replete or sulfate-limited medium provided with 1 mM MT-EtOH versus sulfate-replete medium.

### The Marl gene cluster is required for assimilation of DMS, DMSP, and EMS in R. palustris

To assess if the NFL genes in Mar1, Mar2, or NflDK3 are required for growth on VOSCs, a random barcoded transposon sequencing (RB-TnSeq) library was grown under photoheterotrophic conditions with medium containing 10 mM malate and 1 mM MT-EtOH, and 20 mM acetate with either 1 mM DMS, DMSP, or EMS provided as the sulfur source. As shown in **Fig. 5A**, only insertions in the Mar1 gene cluster result in fitness defects when DMS, DMSP, or EMS is provided as the sulfur source. Interestingly, insertions in the NifX homolog, *nflX* (RPA2355), in Mar2 also lead to fitness defects when DMS and DMSP were provided as a sulfur source (Fig. S2A). Together, this indicates that only the Mar homolog encoded in Mar1 is required for assimilation of DMS, DMSP, and EMS. However, when MT-EtOH was provided as a sulfur source, insertions in genes in Mar1, Mar2, and NflDK3 did not impact fitness. This was unexpected, given that MT EtOH is a substrate for MarHDK in *R. rubrum* (3). We hypothesized that this lack of phenotype may reflect functional redundancy among the NFL enzymes encoded by Mar1, Mar2, or NflDK3 in *R. palustris*.

To further validate the results from the RB-TnSeq and test this hypothesis, we measured the activity of each Mar homolog with each VOSC. To do this, we created *R. palustris* strains that contained a deletion of the NFL genes in Mar1 (ΔMar1), Mar2 (ΔMar2), or NflDK3 (ΔNfl3). *R. palustris* strains containing deletions of two of the three sets of NFL genes were also constructed (ΔMar1Nfl3, ΔMar2Nfl3), as well as a strain lacking all NFL genes in Mar1, Mar2,and NflDK3, which will be referred to as A123. Mar activity was then measured with each of these strains grown with each VOSC used for the RB-TnSeq. Mar activity results in the production of CH_4_ when DMS is provided, C_2_H_4_ when MT-EtOH is provided, and C_2_H_6_ when EMS is provided (3). We measured growth of each mutant and the production of these compounds when their respective VOSC was provided as a sulfur source. As shown in **Fig. 6A** and **Fig. S3A and S3C**, only strains containing a deletion of *marBHDK1* were unable to grow in DMS, EMS, and DMSP. Additionally, CH_4_ production when DMS was provided and C_2_H_6_ production when EMS was provided was dependent on the presence of an intact Mar1 (**Fig. 6B; Fig. S3D**). We also observed CH_4_ production on DMSP, which is consistent with DMSP conversion to DMS and subsequent reduction by MarHDK (**Fig. S3B**). These results indicate that genes in Mar1 encode a methylthio-alkane reductase and has similar activity to the MarHDK from *R. rubrum*. These results also confirm that the other NFL genes in Mar2 and NflDK3 do not appear to play a role in reduction of DMS, DMSP, and EMS.

**Figure 6.**
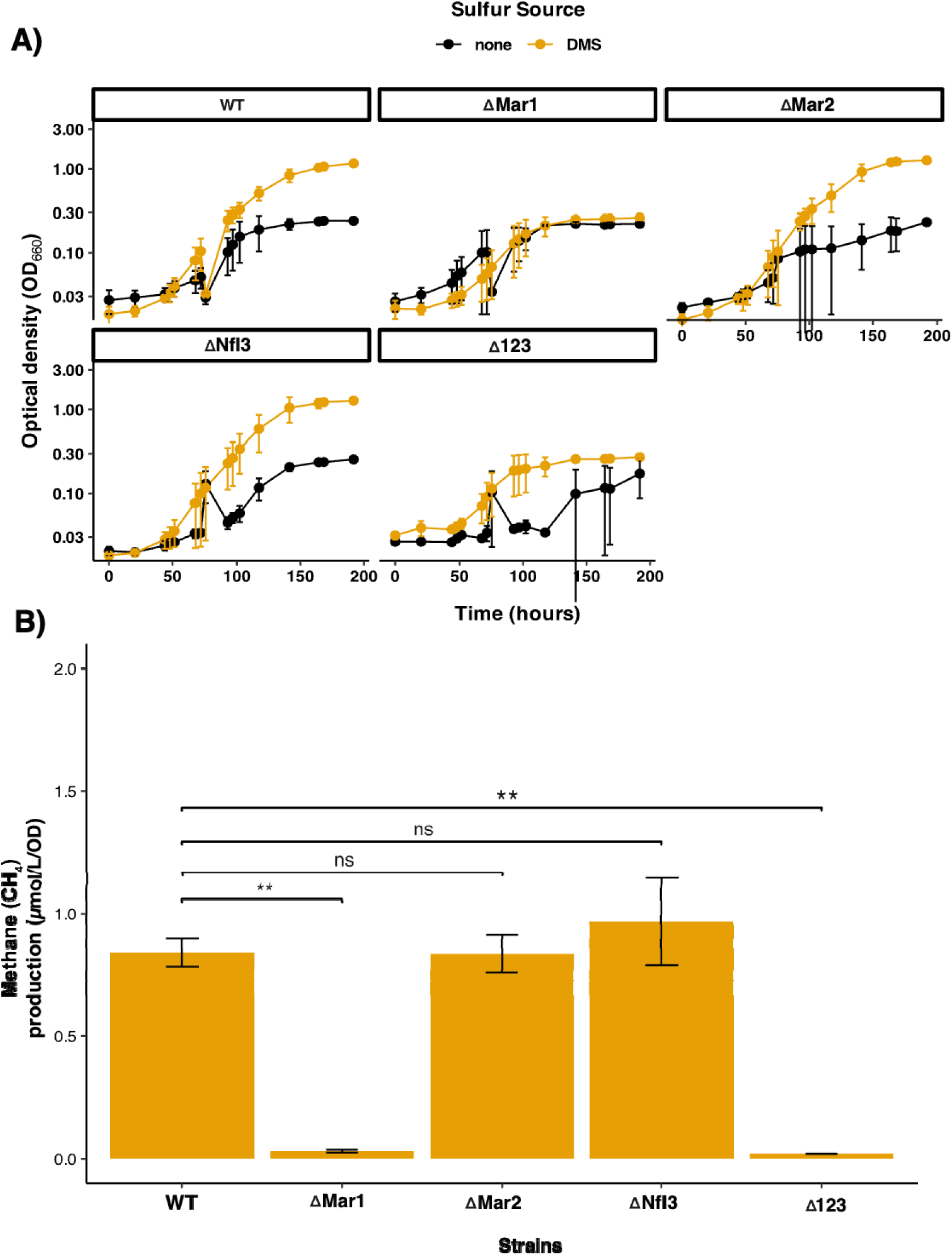
MarBHDK1 (RPA2634-37) is required for the reduction of DMS. **(A)** Growth of *R. palustris* as a measurement of optical density at 660 nm over time in medium with 1 mM DMS as the sulfur source. Cultures were supplied with 20 mM acetate as a carbon source in light. **(B)** Methane (CH_4_) production of wild-type *R. palustris* (WT), *R. palustris* with a deletion in marBHDK1 (ΔMar1), *R. palustris* with a deletion in RPA2347-48 and RPA2353-54 (ΔMar2), *R. palustris* with a deletion in RPA2363-64 (ΔNfl3), and *R. palustris* in which all Mar homologs were deleted (Δ123). Error bars represent the standard deviation of the average of three replicates. Significance was calculated with a t-test. **Significance values:** ns (p > 0.05), * (p ≤ 0.05), ** (p ≤ 0.01), *** (p ≤ 0.001), and **** (p ≤ 0.0001).

One possible explanation for why insertions in any of the NFL genes in Mar1, Mar2, and NflDK3 did not cause a fitness defect in MT-EtOH is that the Mar enzymes are functionally redundant when MT-EtOH is used as a sulfur source. As shown in **Fig. 7A**, *R. palustris* ΔMar1 had only a slightly slower growth rate, and *R. palustris* ΔMar1 cells show reduced but not abolished production of C_2_H_4,_ suggesting one of the other NFL enzymes could be acting on MT-EtOH.

**Figure 7.**
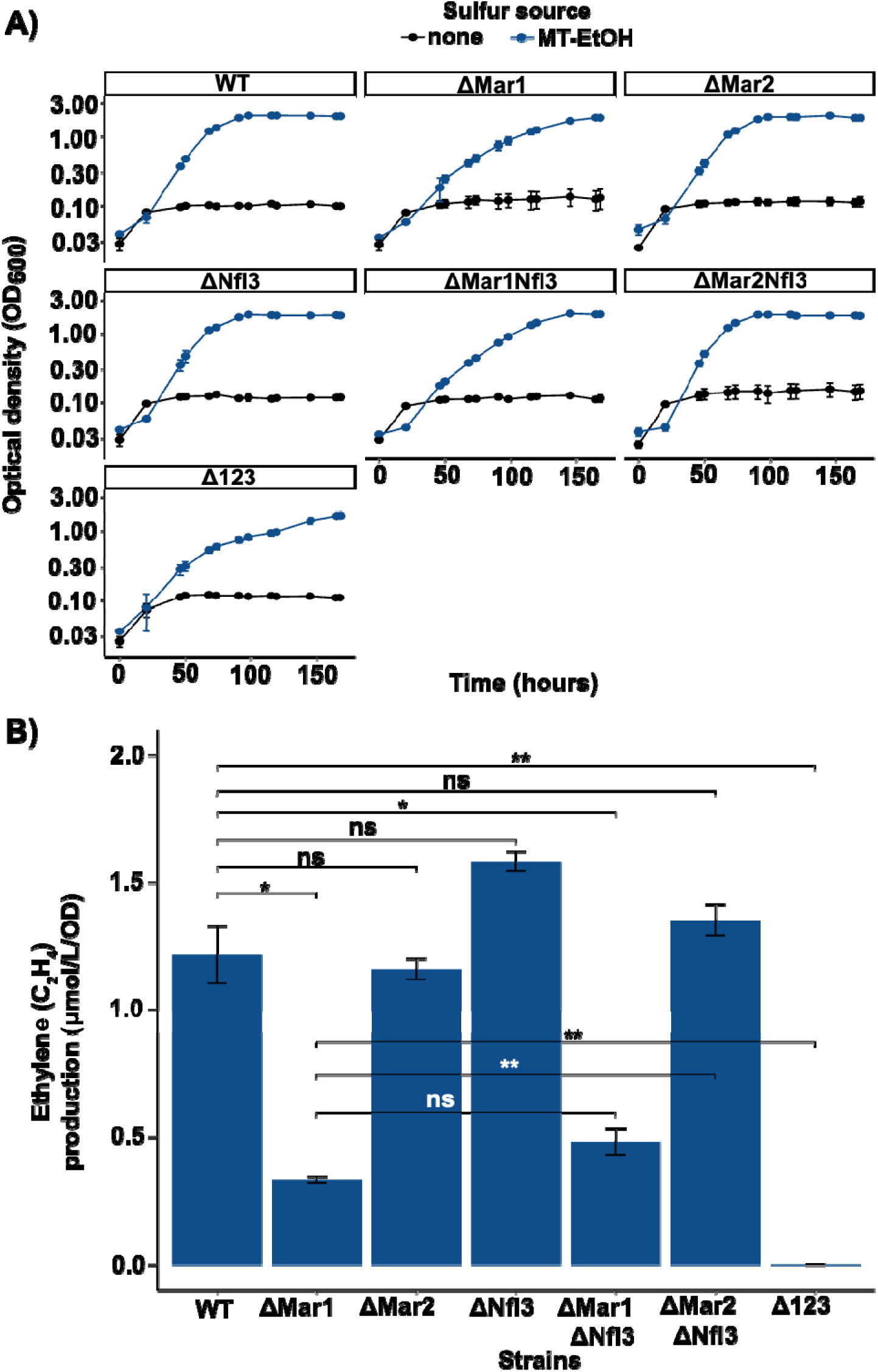
MarBHDK2 (RPA2354-53/RPA2348-47) and MarBHDKI (RPA2634-S7) can reduce MT-EtOH and produce ethylene. **(A)** Growth of *R. palustris* strains in low sulfur media provided with 1 mM of MT-EtOH as the sulfur source and 10 mM malate as a carbon source in the light. Growth assay was done with the following strains: wild-type *R. palustris* (WT), *R. palustris* with a deletion in *marBHDK1* (ΔMar1), *R. palustris* with a deletion in RPA2347-48 and RPA2353-54 (ΔMar2), *R. palustris* with a deletion in RPA2363-64 (ΔNfl3), *R. palustris* strain in which all Mar homologs were deleted (Δ123), a deletion of *marBHDK1* and *nfl3* (ΔMar1ΔNNfl3), and a deletion of *marBHDK2* and *nfl3* (ΔMar2ΔNfl3). **(B)** Ethylene (C_2_H_4_) production of strains grown in A. Error bars represent the standard deviation of the average of three replicates. Significance was calculated with a t-test. **Significance values:** ns (p > 0.05), * (p ≤ 0.05), ** (p ≤ 0.01), *** (p ≤ 0.001), and **** (p ≤ 0.0001).

C_2_H_4_ production was measured in AMar1Nfl3 and AMar2Nfl3 mutants where only Mar1 or Mar2 were present. The presence of Mar2 alone in the ΔMar1 Nfl3 background was sufficient to produce C_2_H_4_ in the presence of MT-EtOH, and only with *R. palustris* A123 was C_2_H_4_ production undetectable (**Fig. 7B**). However, *R. palustris* A123 was still able to grow when MT-EtOH was provided as a sulfur source (**Fig. 7A**), indicating that another Mar-independent pathway for assimilation of MT-EtOH is present, as has been proposed in (2, 22). These results indicate that Mar2 differs in its substrate specificity compared to Mar1 and may be more active with VOSCs that have additional functional groups, like MT-EtOH.

We tested the hypothesis that *R. palustris* Mar2 possesses low activity with MT-ETOH but also had activity with alternative substrates using the recently developed *R. rubrum* complementation system (17). Unlike *R. palustris*, *R. rubrum* only possesses two NFL systems, *marBHDK* and *nflDK*, which may potentially be involved in VOSC metabolism. When these two gene systems are deleted, *R. rubrum* is incapable of metabolizing or growing using DMS, EMS, or MT-EtOH (3). We modified the *R. rubrum* Mar expression plasmid previously constructed to instead express the *marDK1* genes or *marDK2* genes of *R. palustris*. These Mar systems were then expressed in the *R. rubrum* deletion strain and grown in the presence of DMS, EMS, MT-EtOH, or ethanedithiol (EDT) (**Fig. 8**). EDT was chosen given that dithia-alkane species like dithiapentane and dithiahexane are common signaling VOSCS produced by fungi in the rhizosphere where *R. palustris* is found (23, 24). Further, given that in *R. palustris* the native Mar2 system indicated no activity with DMS and EMS, but showed activity with MT-EtOH, this indicated that larger VOSCS may be the preferred substrates. Indeed, when the *R. palustris* Mar1 system was expressed in *R. rubrum*, activity with DMS, EMS, and MT-EtOH, and 10-fold lower activity with EDT was observed (**Fig. 8**). Conversely, the Mar2 system exhibited no activity with DMS or EMS, low activity with MT-EtOH, and higher activity with EDT (**Fig. 8**). The respective activities of Mar1 and Mar2 with DMS, EMS, and MT-EtOH are consistent with those observed when expressed in the native host, *R. palustris* (**Fig. 6B, 7B, and S3**), confirming that Mar2 has residual Mar activity with MT-EtOH, but none with DMS or EMS. Further, these results reveal that Mar2, and to a lesser degree Mar1, is capable of removing the sulfur groups from EDT to produce ethylene, analogous to the reaction with MT-EtOH. As such, Mar2, while retaining some methylthio-alkane reductase activity clearly has evolved in *R. palustris* to catalyze alternative substrates such as dithia-alkanes (i.e. EDT). The substrate profile and substrate specificities for Mar2 are a focus of future studies.

**Figure 8.**
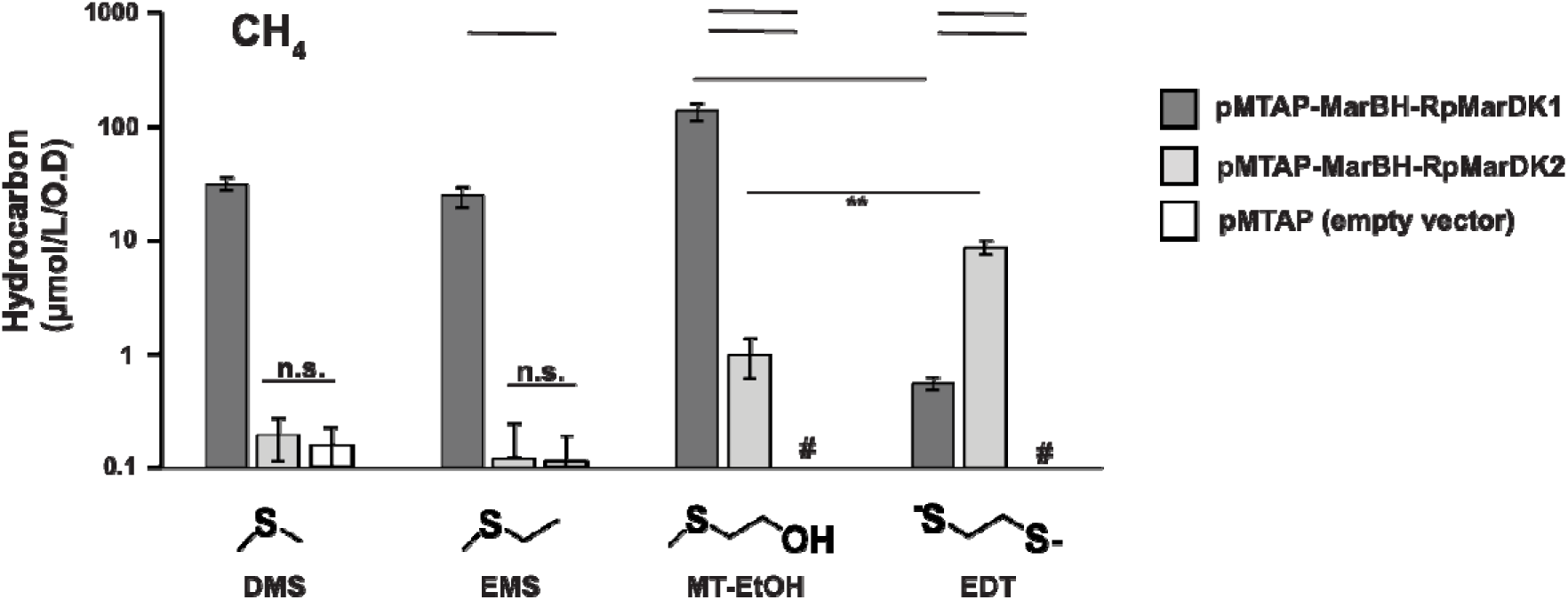
Mar2 of *R. palustris* cleaves mercaptoethanol and ethanethiol. *Rhodospirillum rubrum* Mar deletion strain complemented with no *mar* genes (empty vector), or the *marDK1* and *marDK2* genes of *R. palustris* and grown in the presence of the indicated VOSC. For Mar1, only methane was produced from DMS, ethane from EMS, and ethylene from MT-EtOH or EDT. For Mar2, only ethylene was produced from MT-EtOH and EDT; no activity was observed with DMS and EMS. Hydrocarbon values are for n = 3 independent experiments. #, none detected above limit of 0.1μmol/L/O.D.

To further test Mar activity of Marl and Mar2 from *R. palustris,* hydrogen production w**as** measured. Hydrogen production in the presence and absence of substrate has been detect**ed** with purified MarHDK from *R. rubrum* (18). By measuring hydrogen production, activity of ea**ch** of the Mar homologs in *R. palustris* may be detected even if their substrate is not present. ***R.*** *palustris* CGA009, the strain used in this study, contains a frameshift mutation that results **in** inactivation of its uptake hydrogenase, allowing for the detection of hydrogen produced by M**ar** homologs (25). As shown in **Fig. 9**, wild-type *R. palustris* produced hydrogen when grown w**ith** EMS or MT-EtOH as the sulfur source, and this production was dependent on Mar activity. **As** shown in **Fig. 9A**, most of the hydrogen originated from Mar1 when *R. palustris* was grown with EMS as a sulfur source. However, hydrogen was still detected in ΔMar1, and only A123 show**ed** complete elimination of hydrogen production (**Fig. 9A**), indicating that both Mar1 and Ma**r2** contribute to hydrogen production. In contrast, in the presence of MT-EtOH, ΔMar1 produc**ed** slightly more hydrogen than wild-type *R. palustris,* whereas deletion of ΔMar2 led to a significa**nt** reduction in hydrogen production (**Fig. 9B**). This may reflect that Mar2 exhibits more activity **or** is more highly expressed when grown with MT-EtOH (**Fig. 7B**). Hydrogen production was n**ot** impacted when the NFL genes in NflDK3 were deleted. These results indicate that Mar activity results in hydrogen production *in vivo*, and both Mar1 and Mar2 can produce hydrogen in *R. palustris*.

**Figure 9.**
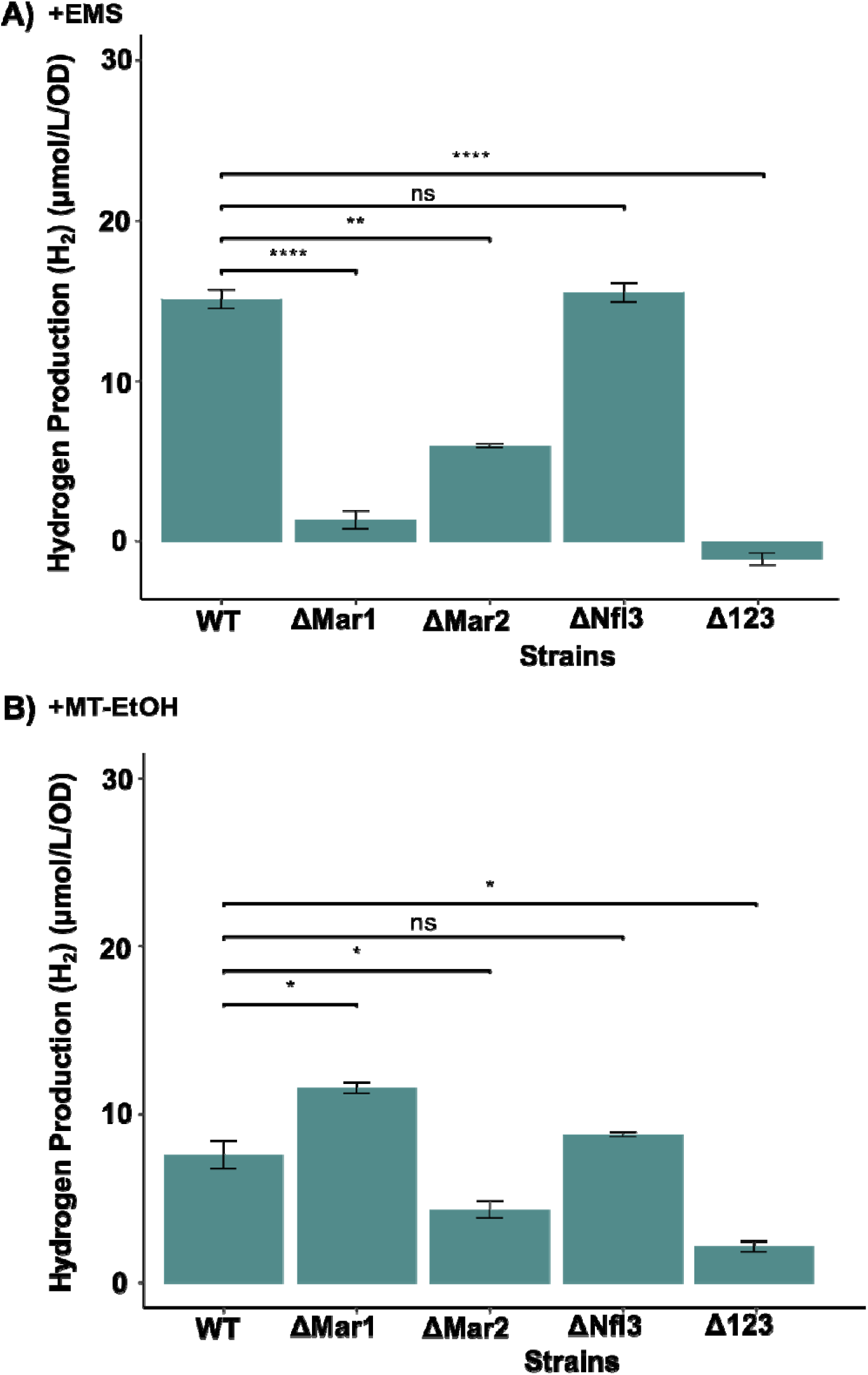
The enzymes encoded by Marl and Mar2 produce H_2_. **(A)** Hydrogen (H_2_) production of wild-type *R. palustris* (WT), a deletion of *marBHDK1* (ΔMar1), a deletion in RPA2354-53 and RPA2348-47 (ΔMar2), a deletion in RPA2363-84 (ΔNfl3), and a strain with a deletion in all Mar homologs (Δ123). Cultures were supplied with 20 mM acetate as a carbon source and 1 mM EMS as sulfur source. **(B)** Hydrogen (H_2_) production of cultures supplemented with 10 mM malate aa a carbon source and 1 mM MT-EtOH as sulfur source. Error bars represent the standard deviation of the mean of the three replicates. Significance was calculated with a t-test. **Significance values:** ns (p > 0.05), * (p ≤ 0.05), ** (p ≤ 0.01), *** (p ≤ 0.001), and **** (p ≤ 0.0001).

### Other bacterial species contain multiple Mar homologs

To evaluate the extent to which individual species harbor multiple Mar homologs, we began by surveying *R. palustris* genomes to identify strains that encode more than one NFL sequence with similarity to MarBHDK. The presence of multiple NFL genes was found in a subset of *R. palustris* strains. We found that *R. palustris* strains YSC-3, PS3, RCB-100, and TIE-1, encode NFL genes similar to those in Mar1, Mar2, and NflDK3 in CGA009 (**Fig. S4**). To determine if other species encode multiple NFL sequences similar to Mar, we identified MarD and MarK homologs in other species. MarD and MarK homologs are mostly found in the phyla Pseudomonadata (Proteobacteria) and Bacillota (Firmicutes). Close homologs of MarDK1 in the Mar1 gene cluster are found in both phyla, but close homologs of MarDK2 (>70% amino acid identity) are restricted to other *Rhodopseudomonas* species, suggesting the NFL genes in Mar2 may have arisen from a gene duplication event in *Rhodopseudomonas.* Close homologs of NflDK3 (>70% amino acid identity) are found in other Alphaproteobacteria isolated from freshwater environments, including *Roseiarcus fermentans, Siculibacillus lacustris, Rhodovastum atsumiense,* and *Blastochloris viridis.* Of these species only *Rhodopseudomonas* species, *Rhodovastum atsumiense,* and *Blastochloris viridis* encode multiple NFL sequences similar to Mar (**Figs. S5 and S6**).

## DISCUSSION

The purpose of this study was to determine the functional roles of the different NFL genes in *R. palustris*. Our findings show that all NFL genes in Mar1, Mar2, and NflDK3 are associated with genes involved in sulfur metabolism and are upregulated under sulfate-limiting conditions. However, despite this shared transcriptional response, only *marHDK1* encoded in Mar1 (RPA2634-37) is required for growth on DMS, DMSP, and EMS. This functional distinction suggests that NFL genes in *R. palustris* are not redundant but instead differ in enzymatic activity or substrate specificity. Consistent with this interpretation, our data indicate that Mar2 acts on MT-EtOH and EDT but not the other VOSCs tested, suggesting it preferentially acts on larger VOSCs that may have different functional groups and are distinct from simpler thioethers like DMS and EMS. Furthermore, restriction of Mar2 to *Rhodopseudomonas* species are consistent with a gene duplication event that may have expanded the capacity of this lineage to use a broader range of organosulfur compounds.

While the location of substrate binding in Mar and homologous systems is unknown, previous structural studies, EPR spectroscopy, mutagenesis studies, and substrate channel modelling by Caver all indicated that the catalysis occurs on the proposed [8Fe-9S-C] metallocofactor (18). In the region surrounding the proposed [8Fe-9S-C] active-site metallocofactor, the *R. rubrum* Mar Cryo-EM structure and the *R. palustris* Mar1 AlphaFold structure show 100% amino acid identity and position (**Fig. S7**). This high similarity between the two homologs is consistent with their respective observed function as methylthio-alkane reductase enzyme for cleavage of MT-EtOH, EMS, and DMS for sulfur acquisitions.

In contrast, amino acid residues in the region of the catalytic metallocofactor are only partially conserved between the *R. rubrum* Mar and the Mar2 of *R. palustris* (**Fig. S8**). Residues MarD H194, W195, S196, F199, F374, and H375 (*R. rubrum* MarD numbering) are conserved between the *R. rubrum* Mar and the *R. palustris* Mar1 and Mar2 systems. These Mar residues are in a similar location to the nitrogenase NifD Q191 and H195 (*A. vinelandii* NifD numbering) for homocitrate, belt sulfur S2B, and substrate coordination. Although MarD residues H194, W195, S196, F199, F374, and H375 may participate in substrate coordination, as suggested by prior mutagenesis studies of W195, their conservation implies that they are unlikely to determine substrate specificity or account for the functional differences observed between Mar and the Mar2 system of *R. palustris*. Interestingly, the main differences between Mar and Mar2 are located along the substrate channel predicted by Caver, as it approaches the catalytic metallocofactor (**Fig. S8**). Specifically, the M70, C73, Y98 and I101 of Mar are replaced with I69, S72, S97, and F101 in Mar2, respectively. Thus, these differences in amino acid species are anticipated to properly coordinate or select for the respective substrates of the Mar systems.

Although the NFL genes found in NflDK3 are expressed in sulfate-limiting conditions or when MT-EtOH is provided as a sulfur source, we did not observe activity with any of the VOSCs tested. Additionally, although we see evidence of hydrogen production from Mar1 and Mar2, a strain that lacks *nflDK3* did not impact hydrogen production on either EMS or MT-EtOH (Fig. 9). This leaves the function of the *nflDK3* still unresolved. *R. rubrum* encodes a second *n*itrogen fixation—*/*ike DK pair (NflDK), but NflDK resides in a different clade within the nitrogenase superfamily (3), indicating *nflDK3* likely represents an uncharacterized but functionally divergent member that is distinct from NflDK from *R. rubrum*.

The presence of two MarB homologs also furthers the complexity of the system in *R. pa/ustris.* Recent structural studies show that the MarDK heterotetramer contains metalloclusters analogous to the nitrogenase P-cluster and an L-like cluster, similar to the FeFeco of the iron-only nitrogenase (17, 18). As discussed above, both MarDK1 and MarDK2 contain the conserved cysteines and histidine residues implicated in P- and L-cluster coordination (Fig. S1C-D), suggesting both MarDK1 and MarDK2 likely bind similar metalloclusters to those found in *R. rubrum* MarHDK. MarB is involved in biosynthesis of the L-like cluster, and NifB can functionally substitute for MarB in *R. rubrum*, reinforcing the role of MarB in maturation of an L-like cluster (17, 18). It is curious then that *R. pa/ustris* encodes multiple NFL sequences similar to MarB. Both MarB1 and MarB2 include key NifB-like motifs and share substantial sequence identity with each other and MarB from *R. rubrum*, which would suggest that both MarB homologs could play a similar role.

Despite this apparent redundancy, only insertions in *marB1* cause fitness defects during growth on DMSP, DMS, and EMS, indicating that MarB2 cannot compensate for MarB1 under these conditions. It is unclear whether this is because each MarB produces a distinct metallocofactor or because there is insufficient expression of *marB2.* Moreover, unlike *R. rubrum, R. palustris* encodes a homolog of NifX that is essential for growth with DMS or DMSP as a sulfur source. NifX binds and stabilizes the cofactor generated by NifB, but it is not essential for nitrogen fixation, implying that Mar in *R. palustris* may involve a more complex system for metallocofactor trafficking (13, 19, 26). Further biochemical and genetic characterization is required to resolve distinctions between multiple NFL genes and to clarify metallocofactor biosynthesis and trafficking in a system with multiple NFL genes.

## Materials and Methods

### Bacteria, culture methods and reagents

*R. palustris* CGA009 and other mutant strains were grown in a defined mineral medium as described in (30). Replacement of ammonium sulfate (NH_4_SO_4_) with ammonium chloride (NH_4_Cl) and a 1 % mineral salts solution containing chloride salts instead of sulfate salts in the defined mineral medium (see Table S1) was used to remove sulfate from medium used in sulfate-limiting conditions. Anoxic media was prepared using an anaerobic chamber with an atmosphere of 98 % N_2_, 2 % H_2_, <10 ppm O_2_ as previously described (31). All *R. palustris* strains were individually grown in 10 mL of mineral minimal media with 20 mM acetate and 0.1 % yeast extract. When the cells reach an optical density at 660nm (OD_660_) of ∼1.0, 100 pL of culture are inoculated in sulfate-limiting medium with 20 mM acetate and 1 mM of either DMS, DMSP, EMS, and 10 mM malate with 1 mM MT-EtOH to acclimate the cultures to the new conditions. Similarly, when cells reach an OD_660_ of ∼0.7-1.0 after acclimation, they were diluted to an OD_660_ of 0.03 in the same medium used for acclimation and OD_660_ was measured over time. All *R. pa/ustris* cultures were grown anoxically in hungate tubes in front of a 60 W incandescent light bulb at 30 °C. *Escherichia co/i* strains were grown in lysogeny broth (LB) at 37 °C. Antibiotics were supplemented as needed. *R. pa/ustris* was given 100 pg/mL of gentamicin or kanamycin, and *E. co/i* was provided with 20 pg/mL of gentamicin. *R. rubrum* strain *kmarBHDK knf/DK,* which is devoid of all NFL genes potentially involved in VOSC metabolism, was transformed with pMTAP empty vector, or the vectors expressing the *R. pa/ustris* Marl or Mar2 systems by conjugative mating as previously described (2).

Transformants were grown anaerobically in sealed culture tubes with Ormerod’s sulfur free malate minimal media with 1 mM ammonium sulfate and without or with 1 mM of a specific VOSC (17). DMS was from Sigma Aldrich, EMS was from TCI America, MT-EtOH and EDT were from Thermo Fisher. Cultures were grown at 30 °C with 2000 lux incandescent illumination until stationary phase was reached.

### Genetic manipulation of R. palustris and R. rubrum

In-frame deletions for each gene of interest were done by creating using pJQ200SK as a suicide vector to introduce a deletion using homologous recombination (26). pJQ200SK was amplified using PrimeSTAR® Max DNA Polymerase (Takara Bio). The pJQ200sk deletion vectors included ∼1 kb of sequence upstream of the start codon and ∼1 kb downstream of the stop codon of the desired deletion. These fragments were amplified using the primers show in Table 1 from *R. pa/ustris* CGA009 genomic DNA with Phusion High Fidelity DNA polymerase (New England Biolabs). Amplified pieces were assembled using homologous recombination in *Escherichia coli* strain DH5- (New England Biolab) as previously described (32). Whole plasmid sequencing was performed by Plasmidsaurus using Oxford Nanopore Technology with custom analysis and annotation. Purified plasmids of confirmed mutants were then transformed in *E. coli* strain S17-1 and subsequently conjugated with *R. palustris.* Gene deletions were confirmed with PCR using primers in Table 1. Whole genome sequencing was also used to assess if there were other mutations, as complementation is not possible in a sulfur limited background since the antibiotic for selection of the complementing vector requires gentamicin sulfate.

**Table 1.**
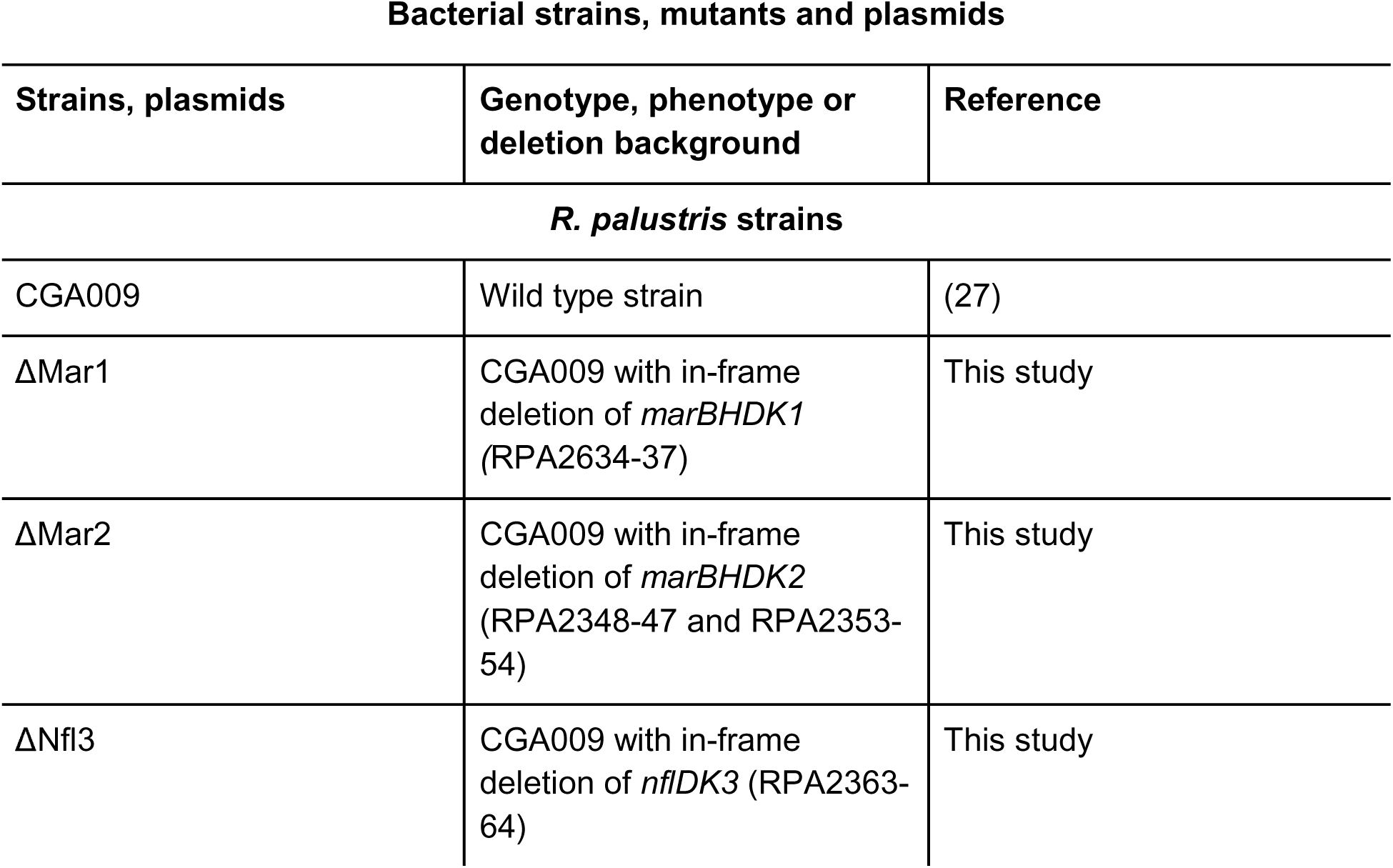
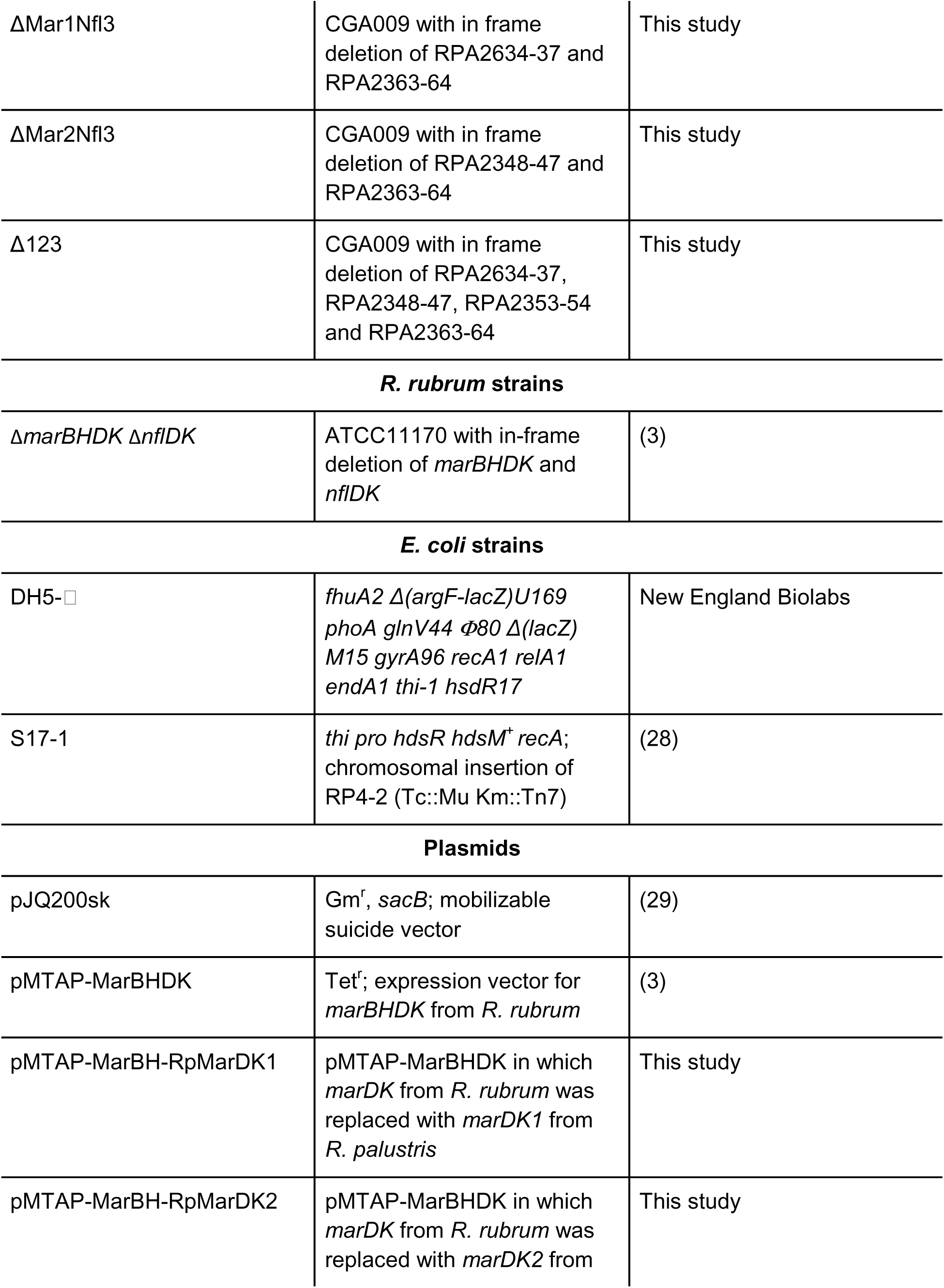
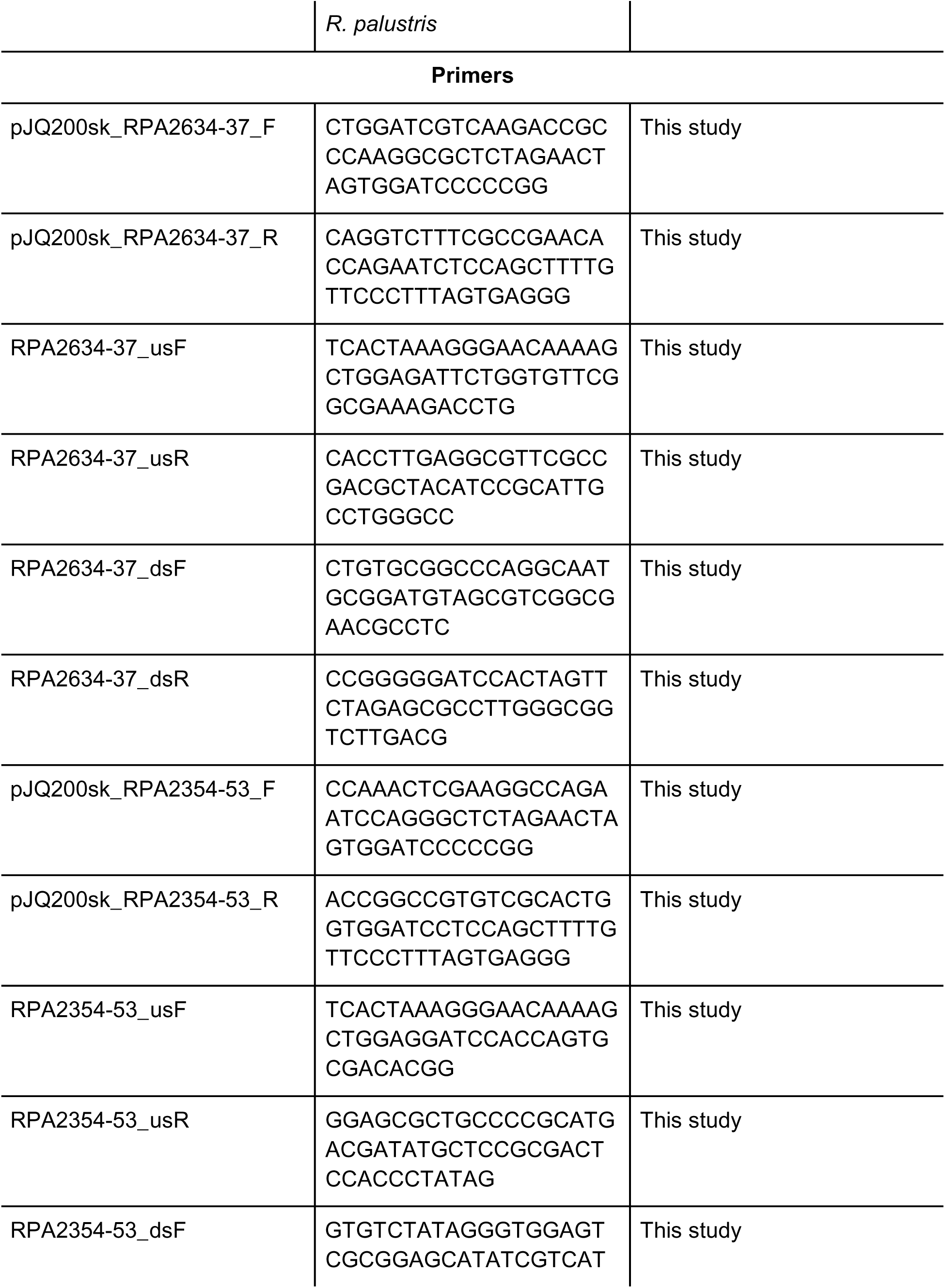
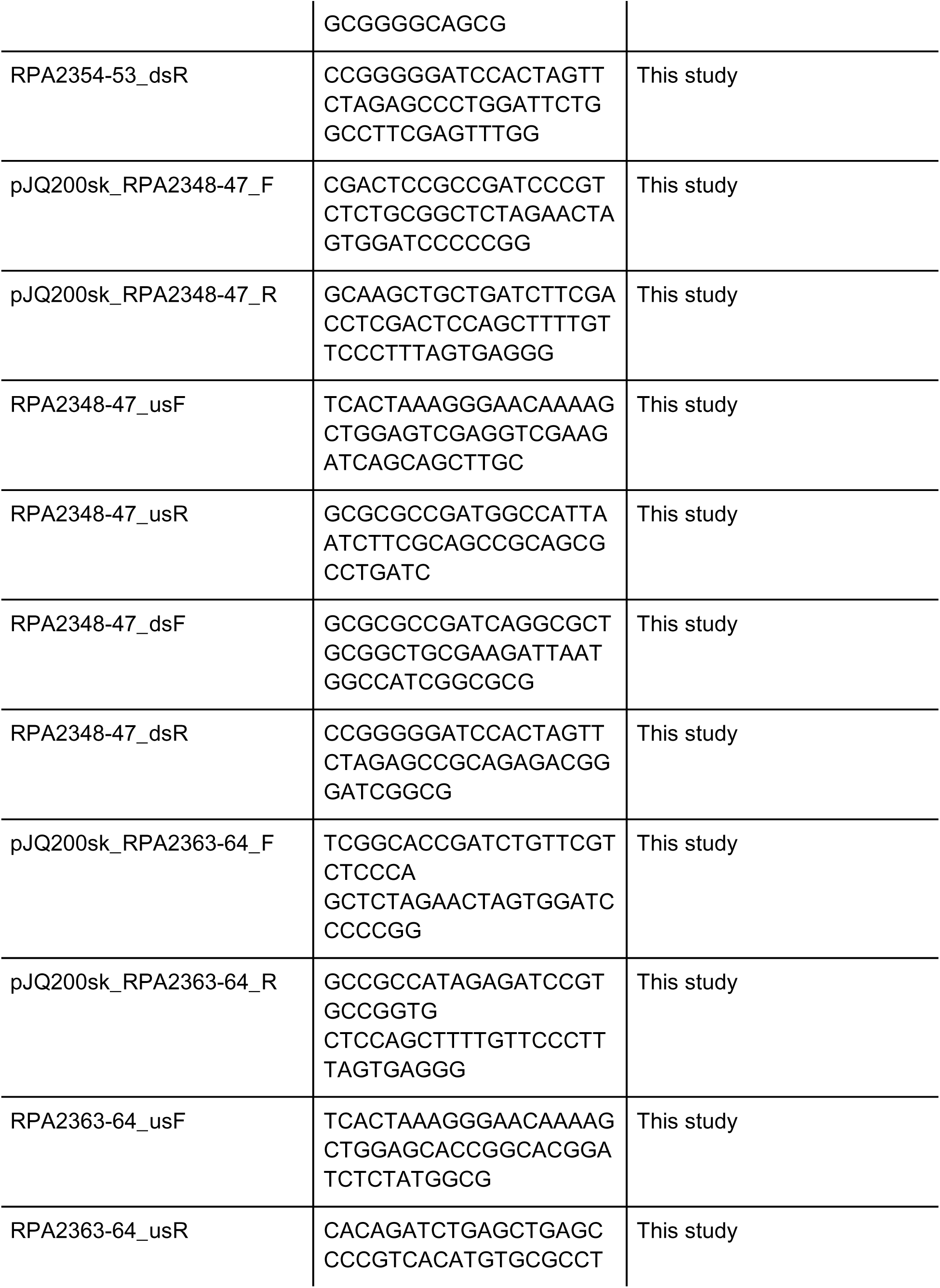
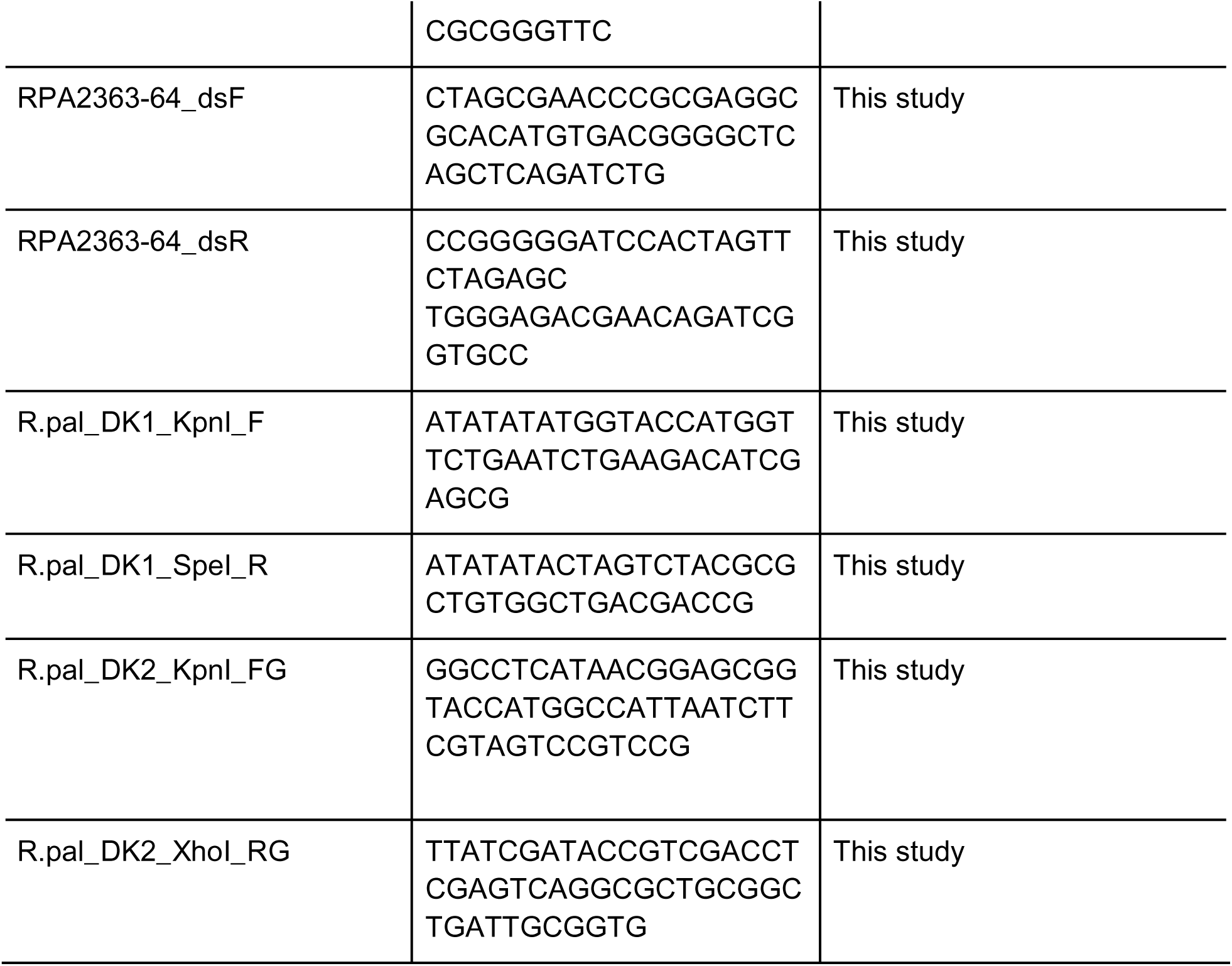
All bacterial strains, plasmids, and primers used in this study.

The *R. palustris marD1* and *marK1* genes were amplified from the genome using *R.pal*_DK1_KpnI_F and *R.pal*_DK1_SpeI_R primers shown in Table 1. Plasmid pMTAP-MarBHDK, which possesses the *R. rubrum marBHDK* genes was digested with Kpnl and Xbal to remove the *R. rubrum marDK* genes (3). The *R. palustris marDK1* PCR fragment was digested with Kpnl and Spel and ligated into the digested plasmid to produce pMTAP-MarBH-RpMarDKI. Similarly, *R. palustris marD2* and *marK2* genes were amplified from the genome using R.pal_DK2_KpnI_FG and R.pal_DK2_XhoI_RG primers shown in Table 1. Plasmid pMTAP-MarBHDK was digested with Kpnl and Xhol to remove the *R. rubrum marDK* genes (3). The *R. palustris marDK2* PCR fragment was inserted into the digested plasmid by Gibson Assembly (New England Biolabs) to produce pMTAP-MarBH-RpMarDK2.

#### Gene neighborhoods

To map out gene neighborhoods, protein sequences (See TableS2) were blasted on IMG Genomes BLAST (https://img.jgi.doe.gov/). With the start and end genome position of the genes in the neighborhood, Gene Cluster Visualizations (v.0.1.10) R 4.3.3 package was used to visualize the clusters.

### Transcriptome sequencing

*R. palustris* CGA009 was grown in mineral medium with 10 mM malate or low sulfur media supplemented with 10 mM malate with and without 1 mM MT-EtOH. Pellets for RNA extraction were harvested at an OD_660_ of 0.2 to 0.4, by incubating 10 mL of cells on ice for 10 minutes, then centrifugation at 15,000 rpm for 10 minutes at 4 °C. Supernatant was removed and cells were flash frozen with liquid nitrogen and stored at −80 °C until further analysis. The pellets were resuspended with 1 mL QIAzol Lysis Buffer and transferred into a 2 mL screw cap tube containing 0.1 mm beads. A Mini-BeadBeater-24 was used to homogenize the samples. The QIAgen miRNeasy Mini Kit (QIAgen, Hilden, Germany) was used to extract total RNA. RNA extraction was followed by a DNAse treatment with Turbo DNAse. Samples were sent to Azenta Life Sciences (https://www.azenta.com/) for Next Generation Sequencing Standard RNA-Seq. Raw data was later trimmed, mapped to the genome of *R. palustris* CGA009, and differential expression analysis on KBase (https://www.kbase.us/). The paired-end reads were trimmed with Trimmomatic (v0.36). The alignment of the trimmed reads to the genome was done using HISAT2 (v2.1.0). The transcripts were assembled from the aligned reads by using StringTie (v.2.1.5). Assembled transcripts were used to do a differential expression analysis matrix with DESeq2 (v.1.20.0). Complete RNA-seq analysis is in Supplemental Dataset 1.

### RB-TnSeq fitness assays

The *R. palustris* CGA009 mutant libraries were thawed and grown in 50 ml of anaerobically prepared mineral minimal medium supplemented with 0.2% casamino acids and 20 mM acetate and 100 pg/ml kanamycin in a 70 mL serum bottle. After cultivation at 30 °C in front of a 60 W lightbulb to a final OD_660_ of ∼2.5, cultures were centrifuged, washed once with PM, and resuspended to an OD_660_ of 3.0. 1 mL aliquots of washed cells were centrifuged, and the resulting cell pellets were stored at −80 °C for later amplicon sequencing and barcode quantification. These aliquots served as the T = 0 samples for RB-TnSeq. The remaining washed cells were diluted to a starting OD_660_ of ∼0.045 in 10 mL of sulfate-limiting medium (Table S1) with different VOSCs provided as a sulfur source and cultivated at 30 °C in the light until stationary phase was reached for each condition. Following completion of growth, cells were harvested for amplicon sequencing and barcode quantification. Fitness scores for all genes in each condition are provided in Supplemental Dataset 2.

### Gas chromatography

Quantification of ethylene, methane, and ethane was done using a Shimadzu GC-2014 with a SUPELCO Analytical 80/100 PORAPAK N 6 ft x 1/8in x 2.1mm SS column. When cells reached mid-log to stationary phase, 100 pL of the headspace was injected into the flame ionization detector (FID) column. Compounds in the headspace are identified based on the retention time and compared with standards of ethylene (Linde), methane (Airgas), and ethane (Airgas). To measure production of H_2_ (Airgas), a 60/80 molecular sieve 5A thermal conductivity detector (TCD) column was used following a similar protocol to that done with the FID column. Samples were measured in three biological replicates and normalized to cell density.

Quantification of hydrocarbons produced by *R. rubrum* strains was performed using a Shimadzu GC-14A with Restek Rt-Alumina BOND/Na_2_SO_4_ column, 30 m, 0.53 mm inner diameter. Gaseous culture headspace after growth experiments was injected at 180□ °C and separated isothermally at 35 □° C. Eluted compounds were detected by flame ionization detector at 180□ °C. From the VOSCs tested, only methane, ethane, and ethylene were observed, and quantified based on methane, ethane and ethylene standard (Schott; Air Liquide).

### Protein alignments and percent identity calculations

*R. palustris* Mar homologs protein sequences (Table S2) were aligned against the protein sequences of *R. rubrum* MarBHDK (Table S2). Sequences were aligned with MUSCLE (33, 34) and regions of conserved residues were annotated with the biopython (v.1.5) package in Python Jupyter notebook (35) following annotations from (3, 15, 18, 20). To calculate percent identity of these protein sequences, the sequences were aligned with the Clustal Omega (36) to calculate a percent identity matrix between sequences.

### Identifying Mar homologs

*R. palustris* Mar protein sequences were separately queried in paperblast (37) to identify the top 100 homologs using MUSCLE to find homologs with ≥ 50% amino acid identity to the sequence queried. The protein sequences of homologs (see Table S3 and S4) were aligned by MUSCLE (33, 34) in R 4.3.3 using the msa package (v1.34.0)(38). A fit maximum likelihood (ML) model was optimized to a Jones-Taylor-Thornton (JTT) model parameters in R 4.3.3 with the phangorn package (v2.12.1) and a neighborhood joining tree was built with the ggtree (v3.17.0) and ggtreeExtra (v.1.12.0) packages. Bootstrap was set for 100 replicates. Percent identity of the homologs was done through Pairwise Alignment with BLOSUM62 through the BioStrings package (v.2.70.3) in R 4.3.3.

### Protein modeling and alignment

AlphaFold 3 with Chai-1 was utilized to predict the structure of MarDK from the *R. palustris* Mar1 and Mar2 systems with coordinated P-cluster and L-cluster (likely mar2 cluster) (39). Metallocofactors were input into Chai-1 as SMILES strings of S_12_[Fe](S([Fe]_22_)[Fe]_3_S_42_)S_3_([Fe]_23_)([Fe]_14_)([Fe]_1_S_22_)[Fe](S_33_)S_1_[Fe]_23_ for the P-cluster and S_12_[Fe](S([Fe]_2_S_23_)[Fe]_33_S_4_)(S_33_([Fe]_567_)([Fe]_12_S_7_)[Fe]_14_S_62_)S[Fe]_3_(S_53_)S_1_[Fe]_32_ for the L-cluster. Bolz-1 and Chai-1 cannot render a true L-cluster from a SMILES string, so an ‘L-cluster’ with a central sulfide was used in the above SMILES string, and then the central sulfide replaced with a carbide in the AlphaFold model. Caver modelling of the *R. rubrum* Mar substrate channel was performed exactly as previously described (18). All structural alignments and visualization were performed using PyMol.

## DATA AVAILABILITY

Transcriptomic data discussed in this paper is available as raw sequencing reads deposited in NCBI’s Gene Expression Omnibus (40, 41) under the GEO Series accession number GSE311060. Gene fitness data is available in the Genome Fitness browser (https://fit.genomics.lbl.gov/cgi-bin/myFrontPage.cgi) under set1S355-362, set1S380-382, set1S384, and set2S385-386 for *R. palustris* CGA009 (42).

## SUPPLEMENTAL MATERIALS

Supplemental Figures and Tables

Table S1 (sulfate-limited medium recipe)

Table S2 (protein accession numbers used to create the percent identity matrix)

Table S3 (protein accession numbers for MarD homologs)

Table S4 (protein accession numbers for MarK homologs)

Supplemental Dataset 1. RNASeq Analysis

Supplemental Dataset 2. VOSC RBTnSeq analysis

## ACKNOWLEDGEMENTS

N.L.M.R was supported by an award from the National Science Foundation Graduate Research Fellowship Program (NSF-GRFP). This work was also supported by award DE-SC0020252 from the U.S. Department of Energy, Office of Science, Basic Energy Sciences, Physical Biosciences program to K.R.F. Substrate determination was supported by the Department of Energy, Office of Science, Biological and Environmental Research EarthShot program (DE-SC0024710 to J.A.N.).

We declare no conflict of interest.

## Notes

### Competing Interest Statement

The authors have declared no competing interest.

### Summary of Updates

Revision includes a title change and supplemental material

## REFERENCES

1. Wetzel RG. 2001. Iron, sulfur and silica cycles, p. 289–330. In Limnology: Lake and River Ecosystems, 3rd ed. Academic Press.

2. North JA, Miller AR, Wildenthal JA, Young SJ, Tabita FR. 2017. Microbial pathway for anaerobic 5-methylthioadenosine metabolism coupled to ethylene formation. Proc Nat/ Acad Sci USA 114:E10455–E10464.

3. North JA, Narrowe AB, Xiong W, Byerly KM, Zhao G, Young SJ, Murali S, Wildenthal JA, Cannon WR, Wrighton KC, Hettich RL, Tabita FR. 2020. A nitrogenase-like enzyme system catalyzes methionine, ethylene, and methane biogenesis. Science 369:1094–1098.

4. Watson SB, Jüttner F. 2017. Malodorous volatile organic sulfur compounds: Sources, sinks and significance in inland waters. Crit Rev Microbio/ 43:210–237.

5. Tallant TC, Krzycki JA. 1997. Methylthiol:coenzyme M methyltransferase from *Methanosarcina barkeri*, an enzyme of methanogenesis from dimethylsulfide and methylmercaptopropionate. J Bacterio/ 179:6902–6911.

6. Tallant TC, Paul L, Krzycki JA. 2001. The MtsA subunit of the methylthiol:coenzyme M methyltransferase of *Methanosarcina barkeri* catalyses both half-reactions of corrinoid-dependent dimethylsulfide: coenzyme M methyl transfer. J Bio/ Chem 276:4485–4493.

7. Lomans BP, van der Drift C, Pol A, Op den Camp HJM. 2002. Microbial cycling of volatile organic sulfur compounds. Ce// Mo/ Life Sci 59:575–588.

8. Tsola SL, Prevodnik AA, Sinclair LF, Sanders IA, Economou CK, Eyice Ö. 2024. *Methanomethylovorans* are the dominant dimethylsulfide-degrading methanogens in gravel and sandy river sediment microcosms. Environ Microbiome 19:51.

9. Dubois M, Van den Broeck L, Inze D. 2018. The pivotal role of ethylene in plant growth. Trends Plant Sci 23:311–323.

10. Schaller GE, Kieber JJ. 2002. Ethylene. Arab Book Am Soc Plant Biol 1:e0071.

11. Smith KA, Russell RS. 1969. Occurrence of ethylene, and its significance, in Anaerobic Soil. Nature 222:769–771.

12. Chatterjee R, Allen RM, Ludden PW, Shah VK. 1997. *In vitro* synthesis of the iron-molybdenum cofactor and maturation of the *nif*-encoded apo-dinitrogenase. J Biol Chem 272:21604–21608.

13. Shah VK, Rangaraj P, Chatterjee R, Allen RM, Roll JT, Roberts GP, Ludden PW. 1999. Requirement of NifX and other *nif* proteins for *in vitro* biosynthesis of the iron-molybdenum cofactor of nitrogenase. J Bacteriol 181:2797–2801.

14. Igarashi RY, Seefeldt LC. 2003. Nitrogen fixation: the mechanism of the Mo-dependent nitrogenase. Crit Rev Biochem Mol Biol 38:351–384.

15. Raymond J, Siefert JL, Staples CR, Blankenship RE. 2004. The natural history of nitrogen fixation. Mol Biol Evol 21:541–554.

16. Harwood CS. 2020. Iron-only and vanadium nitrogenases: fail-safe enzymes or something more? Annu Rev Microbiol 74:247–266.

17. Murali S, Hu G-B, Kreitler DF, Carriedo AA, Lewis LC, Fosu SA, Weaver OG, Buzas EM, Byerly KM, Yoshikuni Y, McSweeney S, Shafaat HS, North JA. 2025. Architecture, catalysis and regulation of methylthio-alkane reductase for bacterial sulfur acquisition from volatile organic compounds. Nat Catal 8:1072–1085.

18. Lago-Maciel A, Soares JC, Zarzycki J, Buchanan CJ, Reif-Trauttmansdorff T, Schmidt FV, Lometto S, Paczia N, Schuller JM, Hansen DF, Heller GT, Prinz S, Hochberg GKA, Pierik AJ, Rebelein JG. 2025. Methylthio-alkane reductases use nitrogenase metalloclusters for carbon-sulfur bond cleavage. Nat Cata/ 8:1086–1099.

19. Moreno-Vivian C, Schmehl M, Masepohl B, Arnold W, Klipp W. 1989. DNA sequence and genetic analysis of the *Rhodobacter capsu/atus nifENX* gene region: Homology between NifX and NifB suggests involvement of NifX in processing of the iron-molybdenum cofactor. Mo/ Gen Genet 216:353–363.

20. Buren S, Jimenez-Vicente E, Echavarri-Erasun C, Rubio LM. 2020. Biosynthesis of nitrogenase cofactors. Chem Rev 120:4921–4968.

21. Kertesz MA. 2000. Riding the sulfur cycle - metabolism of sulfonates and sulfate esters in Gram-negative bacteria. FEMS Microbio/ Rev 24:135–175.

22. Miller AR, North JA, Wildenthal JA, Tabita FR. 2018. Two distinct aerobic methionine salvage pathways generate volatile methanethiol in *Rhodopseudomonas pa/ustris*. mBio 9:e00407–18.

23. Rapior S, Breheret S, Talou T, Bessière J-M. 1997. Volatile flavor constituents of fresh *Marasmius a//iaceus* (Garlic Marasmius). J Agric Food Chem 45:820–825.

24. Tirillini B, Verdelli G, Paolocci F, Ciccioli P, Frattoni M. 2000. The volatile organic compounds from the mycelium of *Tuber borchii Vitt*. Phytochemistry 55:983–985.

25. Rey FE, Harwood CS. 2010. FixK, a global regulator of microaerobic growth, controls photosynthesis in *Rhodopseudomonas pa/ustris*. Mo/ Microbio/ 75:1007–1020.

26. Klassen G, Pedrosa FO, Souza EM, Yates MG, Rigo LU. 1999. Sequencing and functional analysis of the *nifENXorf1orf2* gene cluster of *Herbaspiri//um seropedicae*. FEMS Microbio/ Lett 181:165–170.

27. Larimer FW, Chain P, Hauser L, Lamerdin J, Malfatti S, Do L, Land ML, Pelletier DA, Beatty JT, Lang AS, Tabita FR, Gibson JL, Hanson TE, Bobst C, Torres JLT y, Peres C, Harrison FH, Gibson J, Harwood CS. 2004. Complete genome sequence of the metabolically versatile photosynthetic bacterium *Rhodopseudomonas pa/ustris*. Nat Biotechno/ 22:55–61.

28. Simon R, Priefer U, Puhler A. 1983. A broad host range mobilization system for *In Vivo* genetic engineering: transposon mutagenesis in Gram-negative bacteria. BioTechnol 1:784­791.

29. Quandt J, Hynes MF. 1993. Versatile suicide vectors which allow direct selection for gene replacement in Gram-negative bacteria. Gene 127:15–21.

30. Lewis NM, Sarne A, Fixen KR. 2023. Evolving a new electron transfer pathway for nitrogen fixation uncovers an electron bifurcating-like enzyme involved in anaerobic aromatic compound degradation. mBio 14:e0288122.

31. Fixen KR, Pal Chowdhury N, Martinez-Perez M, Poudel S, Boyd ES, Harwood CS. 2018. The path of electron transfer to nitrogenase in a phototrophic alpha-proteobacterium. Environ Microbiol 20:2500–2508.

32. Kostylev M, Otwell AE, Richardson RE, Suzuki Y. 2015. Cloning should be simple: *Escherichia coli* DH5a-mediated assembly of multiple DNA fragments with short end homologies. PLoS ONE 10:e0137466.

33. Edgar RC. 2004. MUSCLE: a multiple sequence alignment method with reduced time and space complexity. BMC Bioinform 5:113.

34. Edgar RC. 2004. MUSCLE: multiple sequence alignment with high accuracy and high throughput. Nucleic Acids Res 32:1792–1797.

35. Granger BE, Perez F. 2021. Jupyter: thinking and storytelling with code and data. Comput Sci Eng 23:7–14.

36. The UniProt Consortium. 2025. UniProt: the Universal Protein Knowledgebase in 2025. Nucleic Acids Res 53:D609–D617.

37. Price MN, Arkin AP. 2017. PaperBLAST: text mining papers for information about homologs. mSystems 2:10.1128/msystems.00039-17.

38. Bodenhofer U, Bonatesta E, Horejš-Kainrath C, Hochreiter S. 2015. msa: an R package for multiple sequence alignment. Bioinformatics 31:3997–3999.

39. Discovery C, Boitreaud J, Dent J, McPartlon M, Meier J, Reis V, Rogozhnikov A, Wu K. 2024. Chai-1: Decoding the molecular interactions of life. bioRxiv 10.1101/2024.10.10.615955.

40. Edgar R, Domrachev M, Lash AE. 2002. Gene Expression Omnibus: NCBI gene expression and hybridization array data repository. Nucleic Acids Res 30:207–210.

41. Barrett T, Wilhite SE, Ledoux P, Evangelista C, Kim IF, Tomashevsky M, Marshall KA, Phillippy KH, Sherman PM, Holko M, Yefanov A, Lee H, Zhang N, Robertson CL, Serova N, Davis S, Soboleva A. 2013. NCBI GEO: archive for functional genomics data sets—update. Nucleic Acids Res 41:D991–D995.

42. Oda Y, Trotter VV, Hurtado CV, Weinberg BL, Liu H, Biswas A, Chuang Y-C, Haas NW, Heerdink-Santos JPM, Lewis NM, Reyes NM, Mazny BE, Schakel OF, McKinlay JB, Harwood CS, Deutschbauer AM, Fixen KR. 2026. Random barcode transposon-site sequencing of *Rhodopseudomonas palustris* CGA009. Res Sq 10.21203/rs.3.rs-9430391/v1.

