## Supplemental Fig. S1-8 and Supplemental Tables S1-4 for "Expansion of nitrogenase-like enzymes involved in microbial assimilation of volatile organic sulfur compounds"

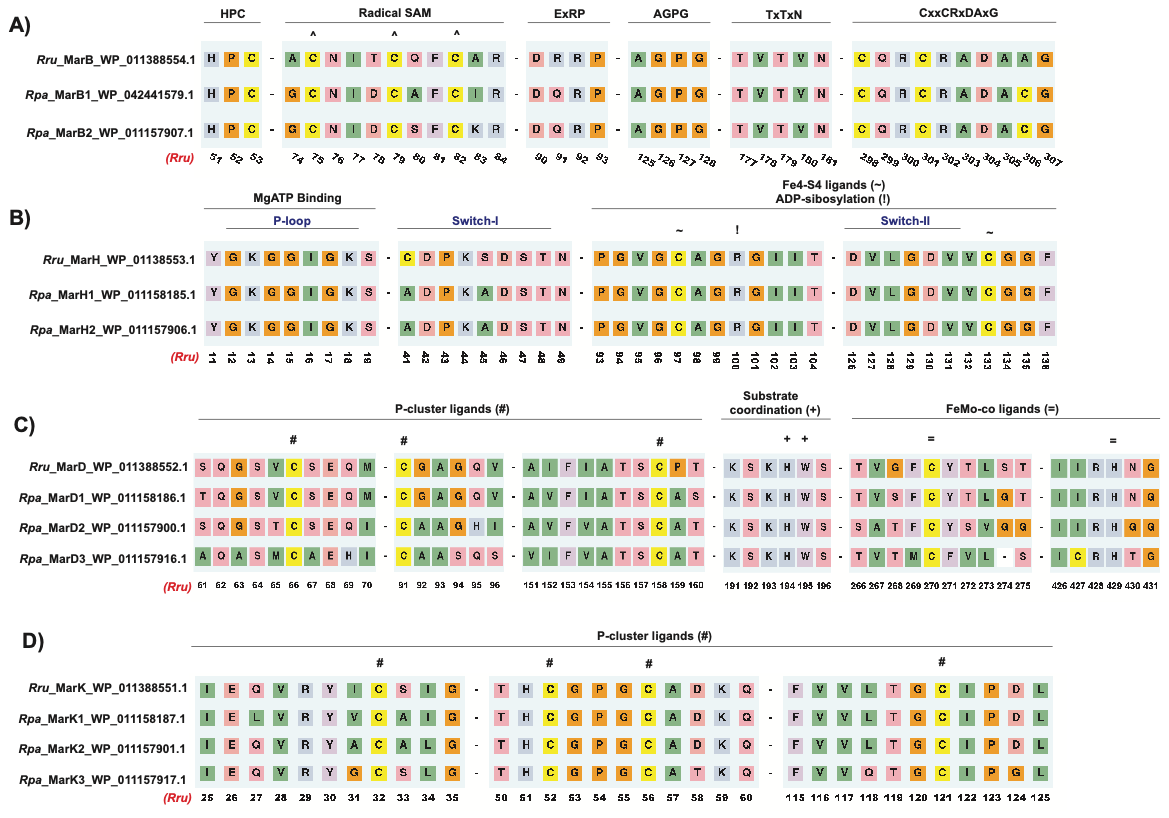

**Fig. S1. MarBHDK sequence alignments.** Numbering is based on the amino acid sequence of the *R. rubrum* Mar protein (1, 3). **(A)** Pairwise alignment of MarB sequences. Sequences share conservation in the radical SAM motif responsible for coordinating 4Fe-4S cluster. **(B)** Pairwise alignment of MarH sequences. Sequences share the MgATP binding motif and the cysteines involved in 4Fe-4S cluster binding. They also share a conserved arginine (shown as “!”) with NifH. This arginine in NifH can be ADP-ribosylated (2). **(C)** Pairwise alignment of MarD sequences. Sequences share conserved residues for P-cluster coordination, L-like ligand, and substrate coordination motifs. **(D)** Pairwise alignment of MarK sequences. Sequences show conservation of the cysteines responsible for coordination of the P-cluster.

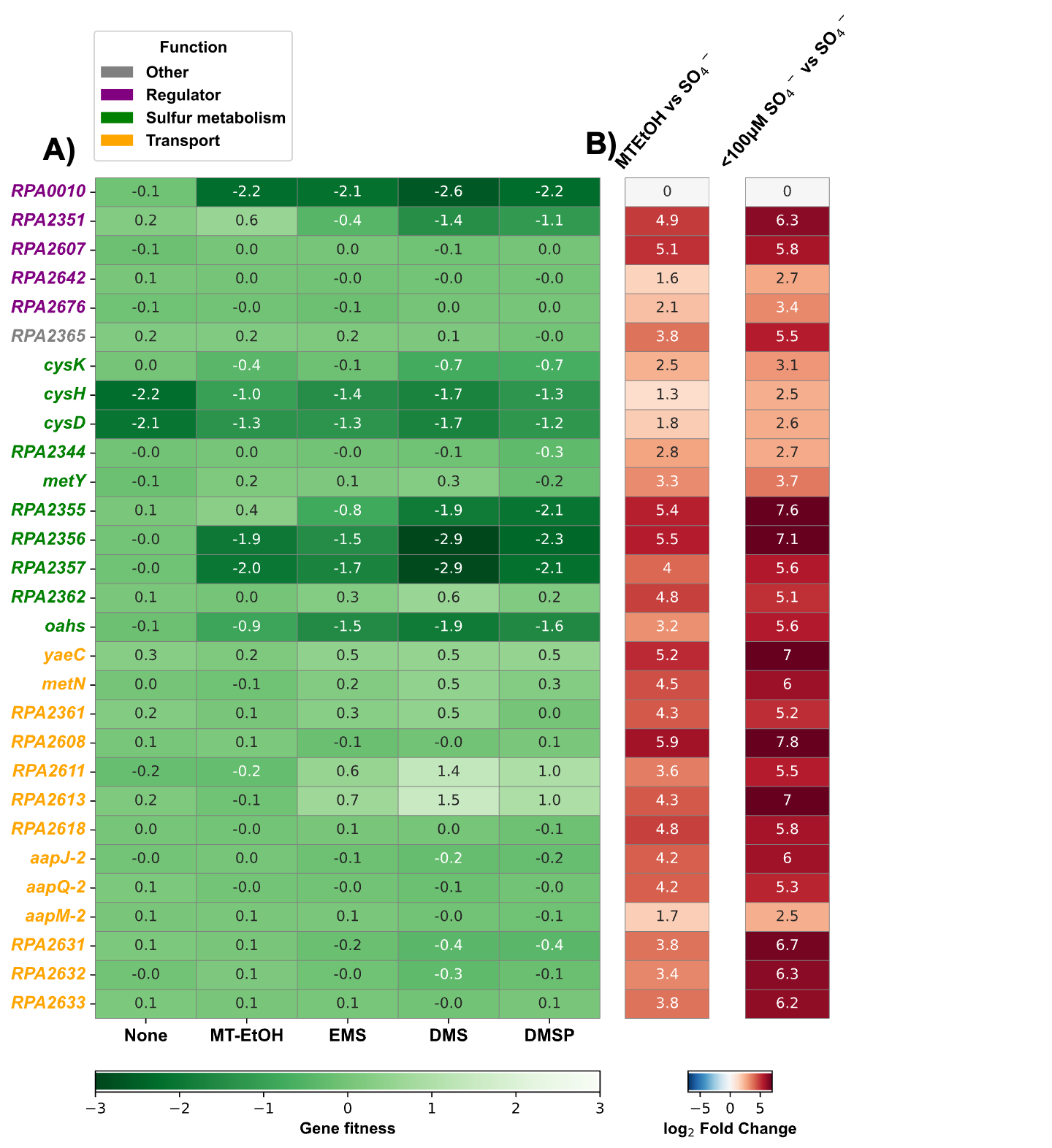

**Fig. S2. Genes involved in sulfur metabolism, sulfur transport and potential sulfur regulation are all upregulated under sulfate limitation, and some genes are required for growth when a VOSC is provided as a sulfur source.** **(A)** RB-TnSeq shows gene fitness calculated from the mean of three samples grown in medium with <100 μM sulfate and no additional sulfur source (None) or provided with 1 mM of VOSC indicated as the sulfur sources. **(B)** The values represent log_2_ fold change in gene expression in cells shifted from sulfate replete (SO_4_^-^) to sulfate-limiting medium or sulfate-limiting medium with 1 mM MT-EtOH provided as the sulfur source.

**Fig. S3. Mar1 is required for growth and Mar activity when DMSP and EMS is provided as the sulfur source.** Growth of *R. palustris* strains in sulfate-limiting medium provided 20 mM acetate and with either 1 mM DMSP **(A)** or 1 mM EMS **(C)**. Mar activity by whole cells was detected by measuring methane production when provided with DMSP **(C)** or ethane production when provided with EMS **(D).** Error bars represent the standard deviation of the average of three replicates. Significance was calculated with a t-test. **Significance values**: ns (p > 0.05), * (p ≤ 0.05), ** (p ≤ 0.01), *** (p ≤ 0.001), and **** (p ≤ 0.0001).

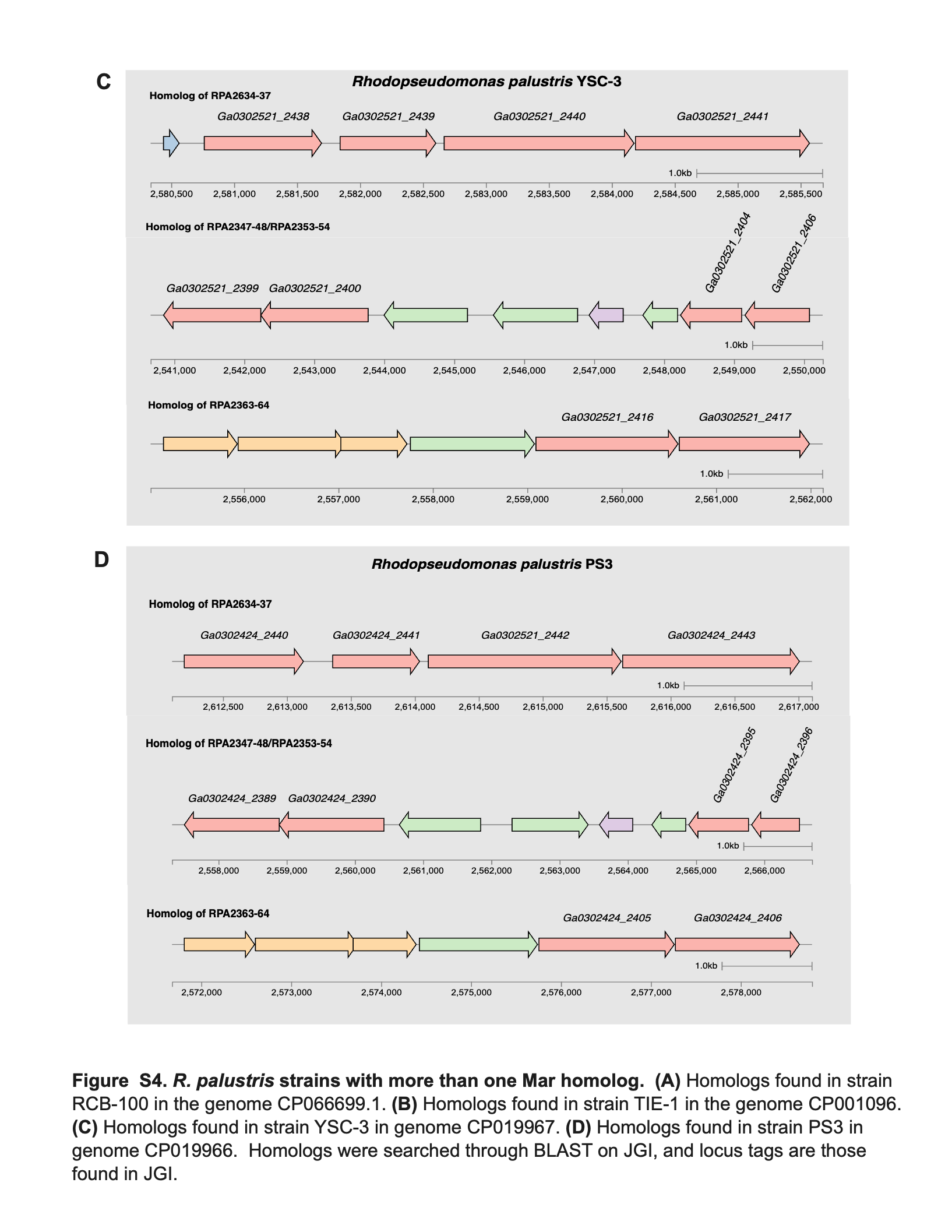
**Fig. S4. Homologous gene clusters of Mar1, Mar2, Nfl3 found in other *R. palustris* strains. (A)** Homologs in strain RCB-100 (Genome: CP066699.1). **(B)** Homologs found in strain TIE-1 (Genome: CP001096). **(C)** Homologs found in strains YSC-3 (Genome: CP019967). **(D)** Homologs found in strain PS3 (Genome: CP0199966). Homologs were searched through BLAST, and locus tags are found in the Integrated Microbial Genomes and Microbiomes from the Joint Genome Instiute (https://img.jgi.doe.gov/).

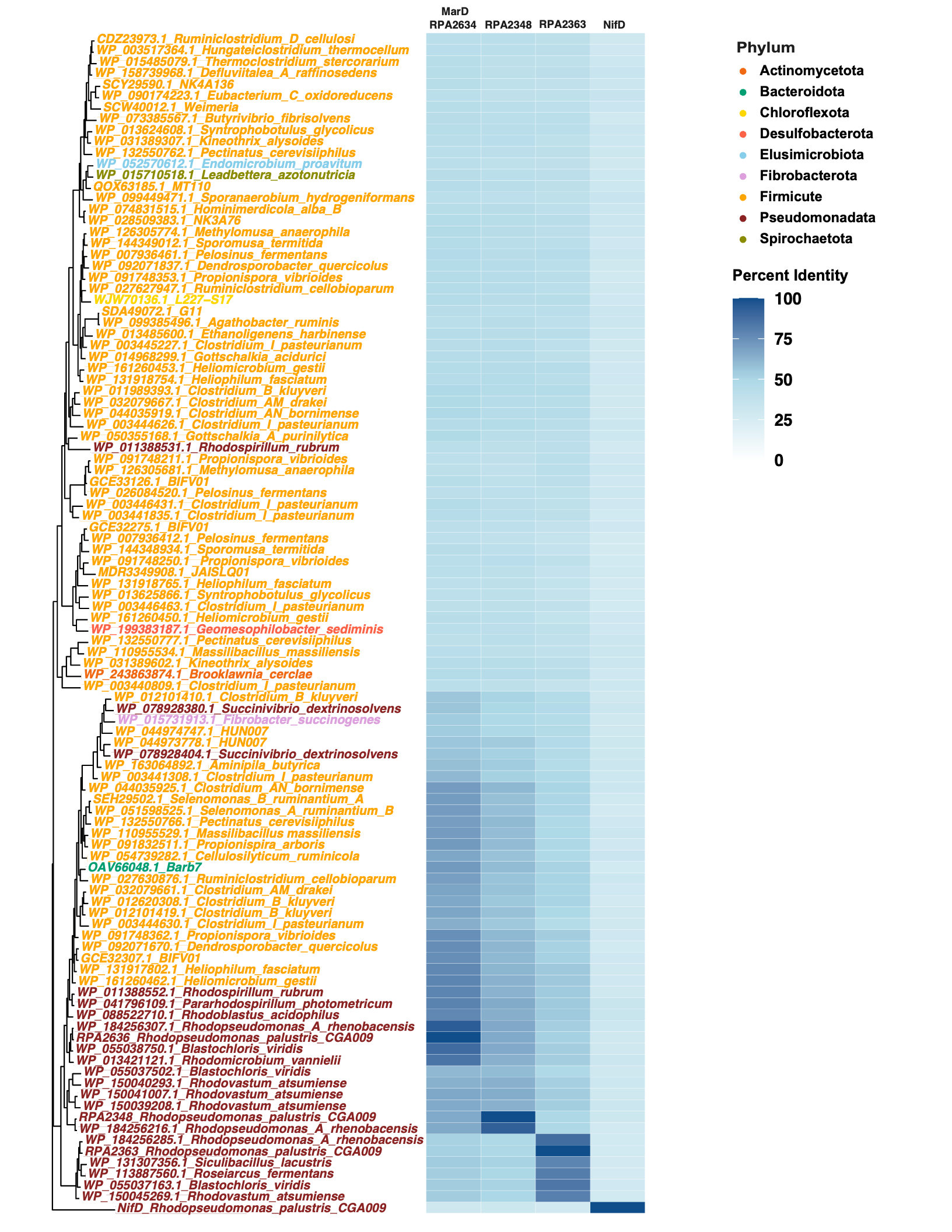
**Fig. S5. Homologs of MarD in different bacterial phyla.** The top 100 blast hits with ≥ 50% amino acid identity to MarD1 (RPA2636), MarD2 (RPA2347) and MarD3 (RPA2363). Blast against *R. palustris’* NifD was also included. Protein sequences of homologs were aligned with MUSCLE and the tree was built as a neighborhood joining tree. The amino acid percent identity to the Mar homologs in *R. palustris* was calculated through BLOSUM62 pairwise alignment.

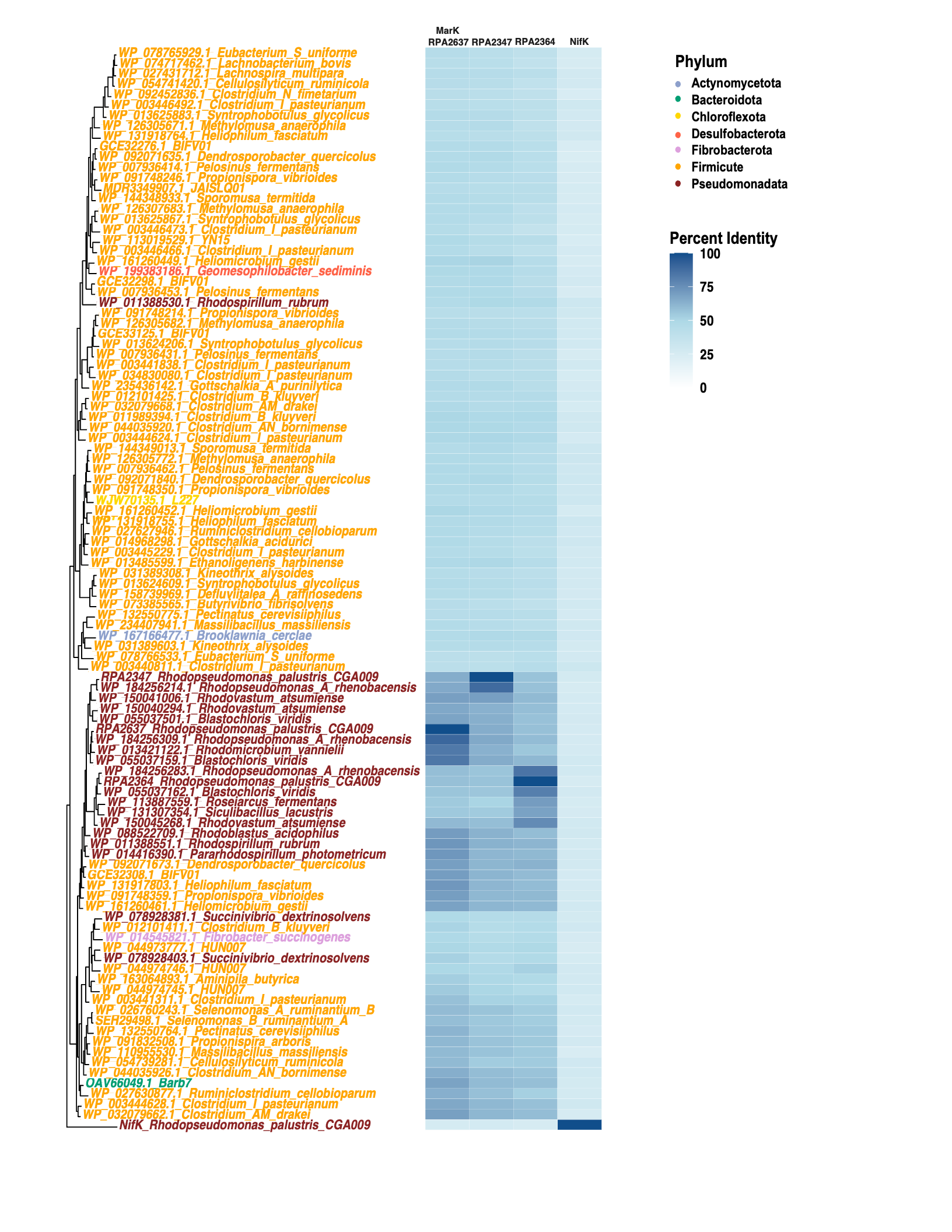

**Fig. S6. Homologs of MarK as found in different bacterial phyla.** The top 100 blast hits with ≥ 50% amino acid to MarK1 (RPA2637), MarK2 (RPA2348), and MarK3 (RPA2364). Blast against *R. palustris’* NifK was done as control. Protein sequences of homologs were aligned with MUSCLE and tree was built as a neighborhood joining tree. The amino acid percent identity to the Mar homologs in *R. palustris* was calculated through BLOSUM62 pairwise alignment.

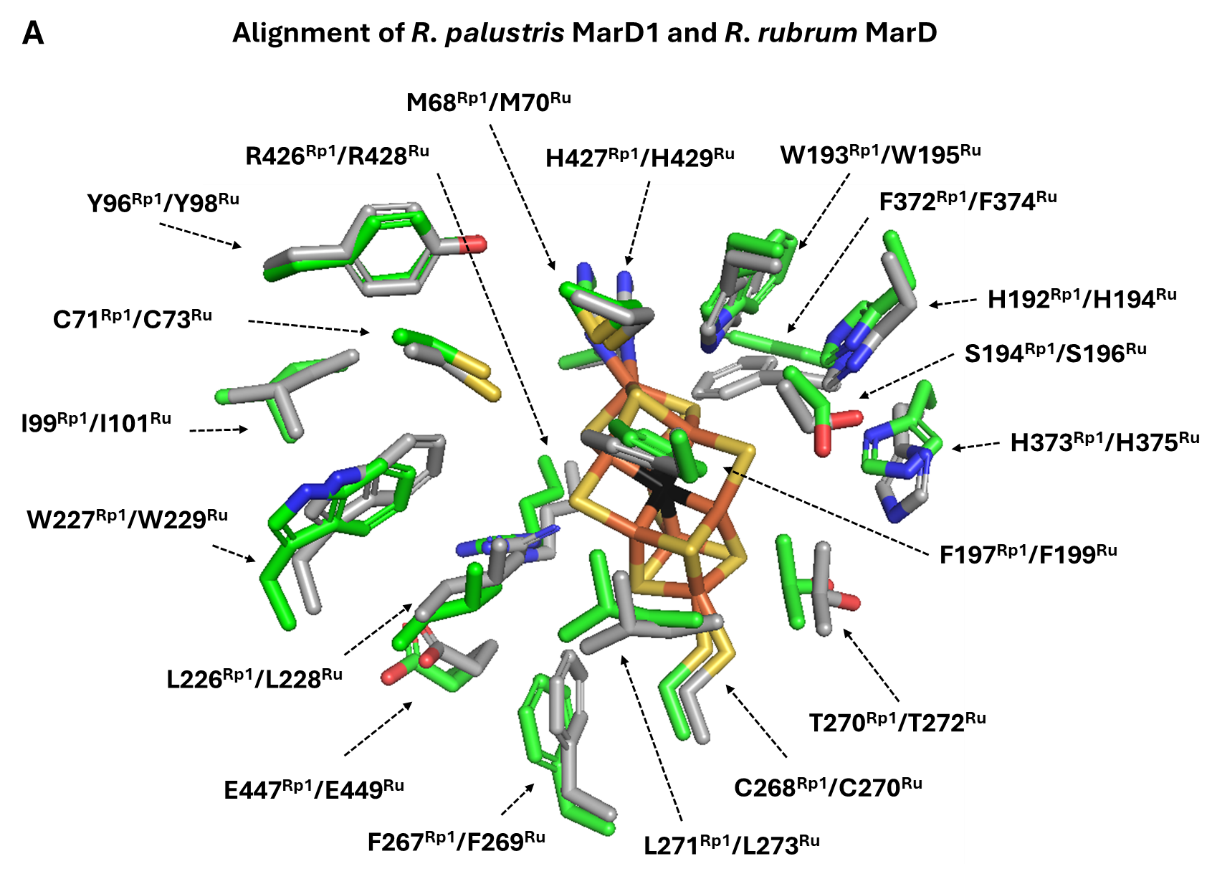

**Figure S7. Alignment of *Rhodospirillum rubrum* MarD structure (9FMG)(2) with an AlphaFold model of *Rhodopseudomonas palustris* MarD of the Mar1 system**. Residues within 20 angstroms of the catalytic mar2 (proposed L-cluster) are displayed. Rp1 superscript indicates the *R. palustris* MarD of the Mar1 system. Ru superscript indicates the *R. rubrum* MarD. There are no differences in residues surrounding the catalytic cofactor between the two Mar enzymes.

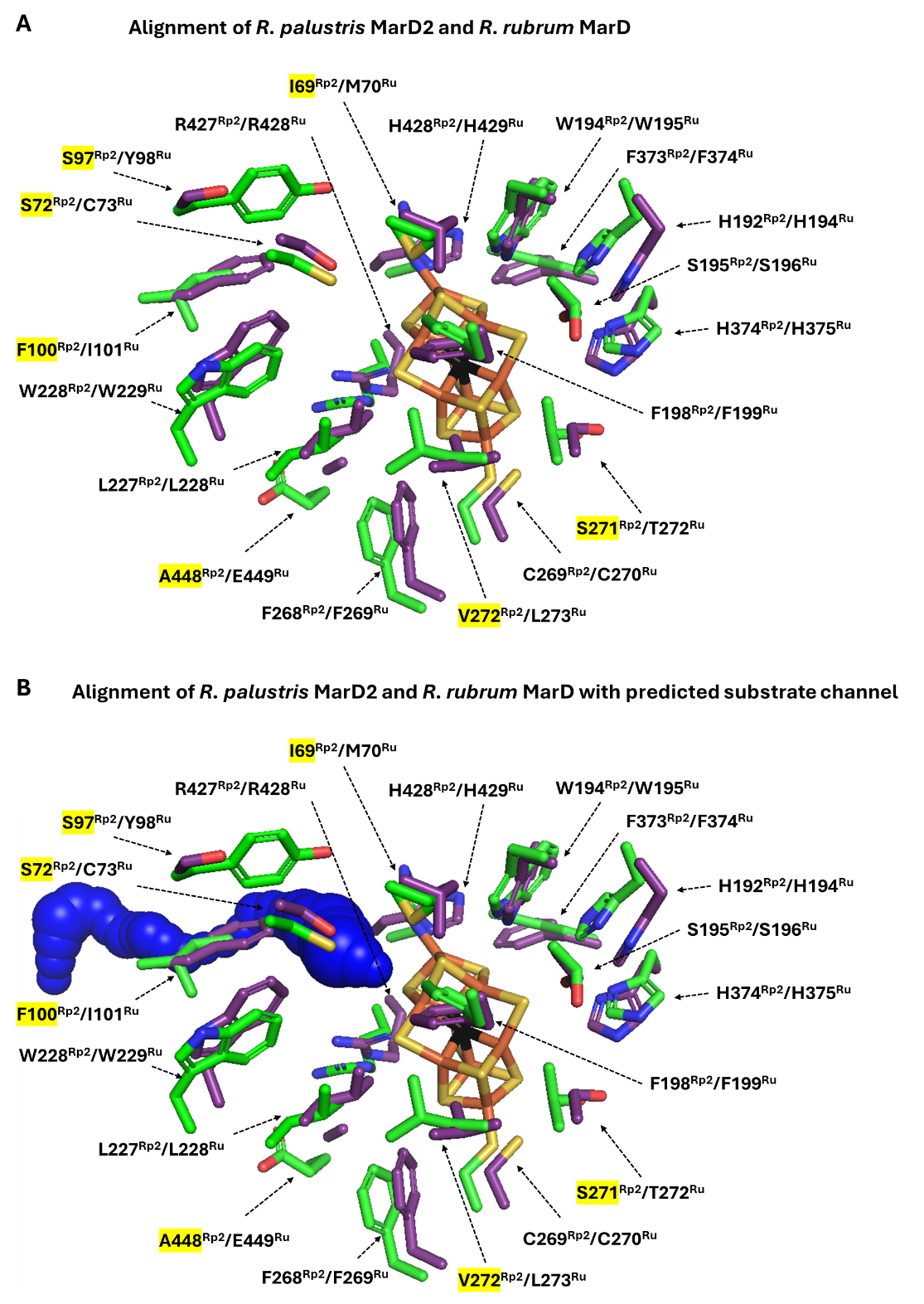

**Figure S8. Alignment of *R. rubrum* MarD structure (9FMG)(2) with an AlphaFold model of *R. palustris* MarD of the Mar2 system**. Alignment without (**A**) and with (**B**) Caver model of *R. rubrum* MarDK substrate channel. Residues within 20 angstroms of the catalytic mar2 (proposed L-cluster) are displayed. Rp2 superscript indicates the *R. palustris* MarD of the Mar2 system. Ru superscript indicates the *R. rubrum* MarD. Differences in residues surrounding the catalytic cofactor between the two Mar enzymes are highlighted.

**Table S1.** Components of sulfate-limiting medium.

| **Sulfate-limiting medium** | **mL/liter of solution** |
| --- | --- |
| 0.5M Na_2_HPO_4_ | 25mL |
| 0.5M KH_2_PO_4_ | 25mL |
| 10% NH_4_Cl | 10mL |
| No sulfur concentrated base | 1mL |
| 2mg/mL p-amino benzoic acid (PABA) | 1mL |
| **No sulfur mineral-salts solution (no sulfur concentrated base)** | **grams/liter of solution** |
| Nitriloacetic acid (NTA-free acid) | 20g |
| MgCl_2_ • 6H_2_O | 48.81g |
| CaCl_2_• 2H_2_O | 6.67g |
| (NH_4_)_6_Mo_7_O_24_• 4H_2_O | 0.0185g |
| FeCl_2_• 4H_2_O | 0.142g |
| No sulfur Metal 44 | 100mL |
| **No sulfur Metal 44 solution** | **grams/liter of solution** |
| EDTA (free acid, not sodium salt) | 2.5g |
| Zinc chloride anhydrous, 98+% | 5.18g |
| Iron (II) chloride • 4H_2_O | 3.58g |
| Manganese chloride • 4H_2_O | 1.78g |
| Copper (II) chloride • 2H_2_O | 0.256g |
| Co(NO_3_)_2_ • 6H_2_O | 0.25g |
| Na_2_B_4_O_7_• 10H_2_O | 0.177g |

**Table S2**. Accession numbers of NifBHDK from *R. palustris* and all MaBHDK homologs from *R. palustris* and *R. rubrum*

| **Species** | **Strain** | **Gene name** | **Protein accession number** |
| --- | --- | --- | --- |
| Rhodopseudomonas palustris | CGA009 | *nifB* | WP_011160162.1 |
| Rhodopseudomonas palustris | CGA009 | *nifH* | WP_011160152.1 |
| Rhodopseudomonas palustris | CGA009 | *nifD* | WP_011160151.1 |
| Rhodopseudomonas palustris | CGA009 | *nifK* | WP_011160150.1 |
| Rhodospirillum rubrum | ATCC 11170 | *marK* | WP_011388551.1 |
| Rhodospirillum rubrum | ATCC 11170 | *marD* | WP_011388552.1 |
| Rhodospirillum rubrum | ATCC 11170 | *marH* | WP_011388553.1 |
| Rhodospirillum rubrum | ATCC 11170 | *marB* | WP_011388554.1 |
| Rhodopseudomonas palustris | CGA009 | *marB1* | WP_042441579.1 |
| Rhodopseudomonas palustris | CGA009 | *marH1* | WP_011158185.1 |
| Rhodopseudomonas palustris | CGA009 | *marD1* | WP_011158186.1 |
| Rhodopseudomonas palustris | CGA009 | *marK1* | WP_011158187.1 |
| Rhodopseudomonas palustris | CGA009 | *marK2* | WP_011157900.1 |
| Rhodopseudomonas palustris | CGA009 | *marD2* | WP_011157901.1 |
| Rhodopseudomonas palustris | CGA009 | *marH2* | WP_011157906.1 |
| Rhodopseudomonas palustris | CGA009 | *marB2* | WP_011157907.1 |
| Rhodopseudomonas palustris | CGA009 | *nflD3* | WP_011157916.1 |
| Rhodopseudomonas palustris | CGA009 | *nflK3* | WP_011157917.1 |

**Table S3.** Protein accession numbers of MarD homologs

| **Species name** | **Protein accession number** |
| --- | --- |
| Hungateiclostridium thermocellum | WP_003517364.1 |
| Ruminiclostridium_D_cellulosi | CDZ23973.1 |
| Defluviitalea A raffinosedens | WP_158739968.1 |
| Thermoclostridium stercorarium | WP_015485079.1 |
| NK4A136 | SCY29590.1 |
| Eubacterium C oxidoreducens | WP_090174223.1 |
| Weimeria | SCW40012.1 |
| Butyrivibrio fibrisolvens | WP_073385567.1 |
| Pectinatus cerevisiiphilus | WP_132550762.1 |
| Kineothrix alysoides | WP_031389307.1 |
| Syntrophobotulus glycolicus | WP_013624608.1 |
| Sporanaerobium hydrogeniformans | WP_099449471.1 |
| NK3A76 | WP_028509383.1 |
| Hominimerdicola alba B | WP_074831515.1 |
| Endomicrobium proavitum | WP_052570612.1 |
| MT110 | QOX63185.1 |
| Leadbettera azotonutricia | WP_015710518.1 |
| G11 | SDA49072.1 |
| Agathobacter ruminis | WP_099385496.1 |
| L227-S17 | WJW70136.1 |
| Ethanoligenens harbinense | WP_013485600.1 |
| Heliomicrobium gestii | WP_161260453.1 |
| Heliophilum fasciatum | WP_131918754.1 |
| Gottschalkia acidurici | WP_014968299.1 |
| Ruminiclostridium cellobioparum | WP_027627947.1 |
| Clostridium I pasteurianum | WP_003445227.1 |
| Dendrosporobacter quercicolus | WP_092071837.1 |
| Pelosinus fermentans | WP_007936461.1 |
| Propionispora vibrioides | WP_091748353.1 |
| Methylomusa anaerophila | WP_126305774.1 |
| Sporomusa termitida | WP_144349012.1 |
| Gottschalkia A purinilytica | WP_050355168.1 |
| Clostridium I pasteurianum | WP_003444626.1 |
| Clostridium AN bornimense | WP_044035919.1 |
| Clostridium B kluyveri | WP_011989393.1 |
| Clostridium AM drakei | WP_032079667.1 |
| Clostridium I pasteurianum | WP_003441835.1 |
| Clostridium I pasteurianum | WP_003446431.1 |
| BIFV01 | GCE33126.1 |
| Pelosinus fermentans | WP_026084520.1 |
| Methylomusa anaerophila | WP_126305681.1 |
| Propionispora vibrioides | WP_091748211.1 |
| Rhodospirillum rubrum | WP_011388531.1 |
| Heliomicrobium gestii | WP_161260450.1 |
| Geomesophilobacter sediminis | WP_199383187.1 |
| Syntrophobotulus glycolicus | WP_013625866.1 |
| Clostridium I pasteurianum | WP_003446463.1 |
| Heliophilum fasciatum | WP_131918765.1 |
| JAISLQ01 | MDR3349908.1 |
| Propionispora vibrioides | WP_091748250.1 |
| Sporomusa termitida | WP_144348934.1 |
| BIFV01 | GCE32275.1 |
| Pelosinus fermentans | WP_007936412.1 |
| MarD3_Rhodopseudomonas_palustris_CGA009 | WP_011157916.1 |
| Rhodopseudomonas A rhenobacensis | WP_184256285.1 |
| Roseiarcus fermentans | WP_113887560.1 |
| Siculibacillus lacustris | WP_131307356.1 |
| Rhodovastum atsumiense | WP_150045269.1 |
| Blastochloris viridis | WP_055037163.1 |
| Blastochloris viridis | WP_055037502.1 |
| Rhodovastum atsumiense | WP_150040293.1 |
| Rhodovastum atsumiense | WP_150041007.1 |
| Rhodovastum atsumiense | WP_150039208.1 |
| MarD2_Rhodopseudomonas_palustris_CGA009 | WP_011157901.1 |
| Rhodopseudomonas A rhenobacensis | WP_184256216.1 |
| Succinivibrio dextrinosolvens | WP_078928380.1 |
| Clostridium B kluyveri | WP_012101410.1 |
| Fibrobacter succinogenes | WP_015731913.1 |
| HUN007 | WP_044974747.1 |
| Aminipila butyrica | WP_163064892.1 |
| HUN007 | WP_044973778.1 |
| Succinivibrio dextrinosolvens | WP_078928404.1 |
| Clostridium I pasteurianum | WP_003441308.1 |
| MarD1_Rhodopseudomonas_palustris_CGA009 | WP_011158186.1 |
| Rhodopseudomonas A rhenobacensis | WP_184256307.1 |
| Blastochloris viridis | WP_055038750.1 |
| Rhodomicrobium vannielii | WP_013421121.1 |
| Pararhodospirillum photometricum | WP_041796109.1 |
| Rhodospirillum rubrum | WP_011388552.1 |
| Rhodoblastus acidophilus | WP_088522710.1 |
| BIFV01 | GCE32307.1 |
| Dendrosporobacter quercicolus | WP_092071670.1 |
| Heliophilum fasciatum | WP_131917802.1 |
| Heliomicrobium gestii | WP_161260462.1 |
| Propionispora vibrioides | WP_091748362.1 |
| Clostridium I pasteurianum | WP_003444630.1 |
| Clostridium B kluyveri | WP_012620308.1 |
| Clostridium B kluyveri | WP_012101419.1 |
| Clostridium AM drakei | WP_032079661.1 |
| Ruminiclostridium cellobioparum | WP_027630876.1 |
| Barb7 | OAV66048.1 |
| Cellulosilyticum ruminicola | WP_054739282.1 |
| Clostridium AN bornimense | WP_044035925.1 |
| Massilibacillus | WP_110955529.1 |
| Propionispira arboris | WP_091832511.1 |
| Pectinatus cerevisiiphilus | WP_132550766.1 |
| Selenomonas A ruminantium B | WP_051598525.1 |
| Selenomonas_B_ruminantium_A | SEH29502.1 |
| Clostridium I pasteurianum | WP_003440809.1 |
| Brooklawnia cerclae | WP_243863874.1 |
| Kineothrix alysoides | WP_031389602.1 |
| Massilibacillus massiliensis | WP_110955534.1 |
| Pectinatus cerevisiiphilus | WP_132550777.1 |
| NifD_Rhodopseudomonas_palustris_CGA009 | WP_011160151.1 |

**Table S4.** Protein accession numbers for MarK homologs

| **Species** | **Protein accession number** |
| --- | --- |
| MarK1_Rhodopseudomonas_palustris_CGA009 | WP_011158187.1 |
| MarK2_Rhodopseudomonas_palustris_CGA009 | WP_011157900.1 |
| MarK3_Rhodopseudomonas_palustris_CGA009 | WP_011157917.1 |
| NifK_Rhodopseudomonas_palustris_CGA009 | WP_011160150.1 |
| Rhodopseudomonas_A_rhenobacensis | WP_184256309.1 |
| Blastochloris_viridis | WP_055037159.1 |
| Rhodomicrobium_vannielii | WP_013421122.1 |
| Heliophilum_fasciatum | WP_131917803.1 |
| Rhodospirillum_rubrum | WP_011388551.1 |
| BIFV01 | GCE32308.1 |
| Propionispora_vibrioides | WP_091748359.1 |
| Dendrosporobacter_quercicolus | WP_092071673.1 |
| Rhodoblastus_acidophilus | WP_088522709.1 |
| Clostridium_AM_drakei | WP_032079662.1 |
| Heliomicrobium_gestii | WP_161260461.1 |
| Rhodovastum_atsumiense | WP_150041006.1 |
| Pararhodospirillum_photometricum | WP_014416390.1 |
| Barb7 | OAV66049.1 |
| Clostridium_I_pasteurianum | WP_003444628.1 |
| Blastochloris_viridis | WP_055037501.1 |
| Rhodovastum_atsumiense | WP_150040294.1 |
| Rhodopseudomonas_A_rhenobacensis | WP_184256214.1 |
| Ruminiclostridium_cellobioparum | WP_027630877.1 |
| Clostridium_AN_bornimense | WP_044035926.1 |
| Pectinatus_cerevisiiphilus | WP_132550764.1 |
| Selenomonas_B_ruminantium_A | SEH29498.1 |
| Propionispira_arboris | WP_091832508.1 |
| Selenomonas_A_ruminantium_B | WP_026760243.1 |
| Cellulosilyticum_ruminicola | WP_054739281.1 |
| Massilibacillus_massiliensis | WP_110955530.1 |
| Clostridium_I_pasteurianum | WP_003441311.1 |
| Rhodovastum_atsumiense | WP_150045268.1 |
| Aminipila_butyrica | WP_163064893.1 |
| Rhodopseudomonas_A_rhenobacensis | WP_184256283.1 |
| HUN007 | WP_044974745.1 |
| Blastochloris_viridis | WP_055037162.1 |
| Siculibacillus_lacustris | WP_131307354.1 |
| Succinivibrio_dextrinosolvens | WP_078928403.1 |
| Clostridium_B_kluyveri | WP_012101411.1 |
| HUN007 | WP_044973777.1 |
| Succinivibrio_dextrinosolvens | WP_078928381.1 |
| Clostridium_I_pasteurianum | WP_003444624.1 |
| Clostridium_B_kluyveri | WP_011989394.1 |
| Clostridium_AN_bornimense | WP_044035920.1 |
| Fibrobacter_succinogenes | WP_014545821.1 |
| Roseiarcus_fermentans | WP_113887559.1 |
| HUN007 | WP_044974746.1 |
| Heliomicrobium_gestii | WP_161260449.1 |
| Clostridium_AM_drakei | WP_032079668.1 |
| Ethanoligenens_harbinense | WP_013485599.1 |
| Clostridium_B_kluyveri | WP_012101425.1 |
| Gottschalkia_A_purinilytica | WP_235436142.1 |
| Geomesophilobacter_sediminis | WP_199383186.1 |
| Pelosinus_fermentans | WP_007936453.1 |
| YN15 | WP_113019529.1 |
| Syntrophobotulus_glycolicus | WP_013625867.1 |
| Methylomusa_anaerophila | WP_126307683.1 |
| Gottschalkia_acidurici | WP_014968298.1 |
| Pelosinus_fermentans | WP_007936462.1 |
| Heliomicrobium_gestii | WP_161260452.1 |
| Methylomusa_anaerophila | WP_126305772.1 |
| Dendrosporobacter_quercicolus | WP_092071840.1 |
| Pelosinus_fermentans | WP_007936414.1 |
| Ruminiclostridium_cellobioparum | WP_027627946.1 |
| Clostridium_I_pasteurianum | WP_003446466.1 |
| BIFV01 | GCE32298.1 |
| Clostridium_I_pasteurianum | WP_034830080.1 |
| Methylomusa_anaerophila | WP_126305682.1 |
| BIFV01 | GCE33125.1 |
| Propionispora_vibrioides | WP_091748350.1 |
| BIFV01 | GCE32276.1 |
| Heliophilum_fasciatum | WP_131918755.1 |
| Syntrophobotulus_glycolicus | WP_013624206.1 |
| Dendrosporobacter_quercicolus | WP_092071635.1 |
| Pelosinus_fermentans | WP_007936431.1 |
| Rhodospirillum_rubrum | WP_011388530.1 |
| Methylomusa_anaerophila | WP_126305671.1 |
| Clostridium_I_pasteurianum | WP_003441838.1 |
| Kineothrix_alysoides | WP_031389603.1 |
| Sporomusa_termitida | WP_144348933.1 |
| Clostridium_I_pasteurianum | WP_003445229.1 |
| Brooklawnia_cerclae | WP_167166477.1 |
| Propionispora_vibrioides | WP_091748246.1 |
| Clostridium_I_pasteurianum | WP_003446473.1 |
| L227-S17 | WJW70135.1 |
| Syntrophobotulus_glycolicus | WP_013624609.1 |
| Pectinatus_cerevisiiphilus | WP_132550775.1 |
| JAISLQ01 | MDR3349907.1 |
| Lachnospira_multipara | WP_027431712.1 |
| Heliophilum_fasciatum | WP_131918764.1 |
| Sporomusa_termitida | WP_144349013.1 |
| Propionispora_vibrioides | WP_091748214.1 |
| Kineothrix_alysoides | WP_031389308.1 |
| Syntrophobotulus_glycolicus | WP_013625883.1 |
| Cellulosilyticum_ruminicola | WP_054741420.1 |
| Eubacterium_S_uniforme | WP_078766533.1 |
| Massilibacillus_massiliensis | WP_234407941.1 |
| Eubacterium_S_uniforme | WP_078765929.1 |
| Clostridium_I_pasteurianum | WP_003440811.1 |
| Lachnobacterium_bovis | WP_074717462.1 |
| Clostridium_N_fimetarium | WP_092452836.1 |
| Clostridium_I_pasteurianum | WP_003446492.1 |
| Defluviitalea_A_raffinosedens | WP_158739969.1 |
| Butyrivibrio_fibrisolvens | WP_073385565.1 |

**Supplemental References**

1.	Murali S, Hu G-B, Kreitler DF, Carriedo AA, Lewis LC, Fosu SA, Weaver OG, Buzas EM, Byerly KM, Yoshikuni Y, McSweeney S, Shafaat HS, North JA. 2025. Architecture, catalysis and regulation of methylthio-alkane reductase for bacterial sulfur acquisition from volatile organic compounds. *Nat Catal* 8:1072–1085.

2.	Lago-Maciel A, Soares JC, Zarzycki J, Buchanan CJ, Reif-Trauttmansdorff T, Schmidt FV, Lometto S, Paczia N, Schuller JM, Hansen DF, Heller GT, Prinz S, Hochberg GKA, Pierik AJ, Rebelein JG. 2025. Methylthio-alkane reductases use nitrogenase metalloclusters for carbon–sulfur bond cleavage. *Nat Catal* 8:1086–1099.

3.	North JA, Narrowe AB, Xiong W, Byerly KM, Zhao G, Young SJ, Murali S, Wildenthal JA, Cannon WR, Wrighton KC, Hettich RL, Tabita FR. 2020. A nitrogenase-like enzyme system catalyzes methionine, ethylene, and methane biogenesis. *Science* 369:1094–1098.
